# Kidney-specific *Wdr72* deletion leads to incomplete distal renal tubular acidosis through impaired V-ATPase B1 subunit localization

**DOI:** 10.1101/2025.10.29.684588

**Authors:** Amr Al-Shebel, Phoebe Mossmann, Sarah Wendlinger, Tilman Breiderhoff, Michael M. Kaminski, Dominik Müller, Philip Bufler, Verena Klämbt

**Author notes:** Corresponding author: Dr. med. Verena Klämbt, Augustenburger Platz 1, 13353 Berlin.

## Abstract

**Background:** Distal renal tubular acidosis (dRTA) is a rare kidney disorder characterized by impaired urinary acidification due to defective proton secretion in type A intercalated cells of the collecting duct. Recently, pathogenic variants in the human gene encoding the *WD Repeat Domain 72 protein* (*WDR72*) have been reported in patients with dRTA, yet the physiological role of WDR72 in the kidney remains unknown.

**Methods:** To elucidate the renal function of Wdr72, we generated a kidney-specific knockout mouse model (*Wdr72^fl/fl^;Pax8-Cre^+^*) and assessed acid–base homeostasis under baseline, acute, and chronic acid loading.

**Results:** *Wdr72^fl/fl^;Pax8-Cre^+^* mice displayed persistently elevated urinary pH, reduced titratable acid and net acid excretion under basal and acid-loaded conditions, consistent with incomplete dRTA. While the systemic pH remained unchanged compared to controls under standard diet, chronic acid load led to mild hyperchloremic, hypokalemic metabolic acidosis. Notably, urinary NH₄⁺ excretion was increased upon acid loading accompanied by upregulation of key ammoniagenesis enzymes, which was detected even under basal conditions, consistent with a compensatory activation of proximal tubular acid excretion pathways. The total and membranous abundance of the V-ATPase B1 subunit decreased markedly within the kidney, despite unchanged transcript levels, suggesting a defect in V-ATPase trafficking or assembly. In addition, morphometric analyses revealed an increased proportion of type A intercalating cells that failed to expand upon acid loading, indicating defective adaptive plasticity.

**Conclusions:** Kidney-specific *Wdr72* deletion impairs distal urinary acidification through reduced V-ATPase abundance and membranous targeting, altered intercalated cell morphology, and limited adaptive remodeling, resulting in incomplete dRTA. Upregulation of renal ammoniagenesis partially compensates the acidification defect. These findings highlight WDR72 as a key regulator of distal nephron acid–base homeostasis and offer mechanistic insight into WDR72-associated dRTA.

**Key Points:**

- Kidney-specific deletion of *Wdr72* reduced Atp6v1b1 membranous localization in intercalated cells.
- Kidney-specific *Wdr72* knockout altered intercalated cell morphology, and limited their adaptive remodeling.
- The lack of the renal Wdr72 resulted in incomplete dRTA, compensated partially by elevated ammoniagenesis.

## Introduction

Primary distal renal tubular acidosis (dRTA) is a rare inherited disorder of renal acid–base regulation caused by defective proton secretion in the intercalated cells (ICs) of the collecting duct (CD).^1^ Affected individuals typically present in early childhood with failure to thrive, polyuria and hyperchloremic, hypokalemic, metabolic acidosis, which often leads to nephrocalcinosis, nephrolithiasis, and bone demineralization.^1^ Despite lifelong alkali supplementation, many patients develop chronic kidney disease after puberty.^2–4^ In contrast, incomplete forms of dRTA manifest with normal serum pH but defective urinary acidification upon acid challenge.^5^

In pediatric populations, dRTA most commonly results from pathogenic variants in genes encoding key transport proteins of ICs, including *ATP6V0A4*, *ATP6V1B1* (encoding subunits of the vacuolar H⁺-ATPase (V-ATPases))^6–8^, and *SLC4A1* (encoding anion exchanger AE1).^9^ Recently, rare bi-allelic variants in *WDR72* have been identified in patients with dRTA and amelogenesis imperfecta (AI)^10–16^, expanding the genetic spectrum of the disease. However, the physiological role of WDR72 in the kidney and its contribution to dRTA pathogenesis remain largely unknown.

The kidney maintains systemic acid–base balance through different complementary processes: reabsorption of filtered bicarbonate, excretion of acid as ammonium and titratable acids, and generation of new bicarbonate and ammonia via renal ammoniagenesis. Within the CD, acid–base regulation is fine-tuned by distinct specialized IC subtypes according to physiological cues. At least two major IC subtypes are recognized in the distal nephron: type A and type B ICs.^17^ In addition, a third population, referred to as type non-A/non-B ICs, has been proposed, though it is comparatively poorly understood^18^. Type A ICs secrete protons via apical V-ATPases and export newly generated bicarbonate across the basolateral membrane through the chloride/bicarbonate exchanger AE1, particularly under acidotic conditions.^17^ In contrast, non-A ICs (including type B and non-A/non-B ICs) express the chloride/bicarbonate exchanger pendrin at the apical membrane and localize the V-ATPase either basolaterally or apically. Type B ICs mediate bicarbonate secretion and are primarily engaged during systemic alkalosis.^17,19^

V-ATPases are large, multi-subunit proton pumps composed of a membrane-integrated V₀ domain and a cytosolic V₁ domain, whose reversible assembly at the apical membrane is required for active proton transport.^19,20^ Their localization and activity can be dynamically regulated through vesicular trafficking and reversible assembly of the V₀/V₁ complexes in response to systemic acid–base cues.^19,20^ Rabconnectin-3 complexes, consisting of an α-subunit (DMXL1 or DMXL2) and a β-subunit (WDR7), are key modulators of this process, facilitating V-ATPase assembly and stabilization, analogous to the RAVE complex in yeast.^19^

WDR72 is a large cytoplasmic protein containing seven N-terminal WD40 repeat domains predicted to form a β-propeller structure that mediates protein–protein interactions.^21^ It shares ∼37% sequence identity and 58% similarity across the N-terminal region with its homologue WDR7,^21^ the Rabconnectin-3 β-subunit.^21^ *Wdr72* is highly expressed in ICs of the CD.^22^ *Wdr72*-deficient mice have been studied primarily in the context of hypomaturation-type AI, developing poorly mineralized enamel due to defects in the maturation phase,^23,24^ potentially resulting from disrupted vesicle trafficking and altered pH homeostasis in ameloblasts, which are mediated by V⁺-ATPase dependent mechanisms.^23^ Although the precise function of WDR72 remains unknown, its domain architecture, sequence homology to WDR7, and reported role in enamel maturation suggest that WDR72 may regulate vesicle trafficking or assembly, and V-ATPase dependent acidification in epithelial cells. These observations may indicate that WDR72 contributes to the regulation of V-ATPase localization and function in renal ICs, thereby influencing distal nephron acid–base transport.

To investigate the physiological role of Wdr72 in distal nephron acid–base handling, we generated a kidney-specific *Wdr72* knockout mouse model. *Wdr72*-deficient mice displayed features of incomplete dRTA, including impaired urinary acidification, altered IC structure and adaptability, accompanied by compensatory mechanisms. These results reveal Wdr72 as a novel modulator of acid–base regulation in the distal nephron.

## Methods

### Animals

All animal procedures were conducted in compliance with institutional and national animal welfare regulations and approved under license number G 066/22 by the Landesamt für Gesundheit und Soziales (LAGeSo), Berlin, Germany. Mice were maintained under standardized conditions (12-hour light/dark cycle, 22–24 °C ambient temperature, 55 ± 15 % relative humidity) with ad libitum access to standard chow and water. Wdr72^tm1a(KOMP)Wtsi^ mice were obtained from the Knockout Mouse Project (KOMP) repository.^25^ To generate kidney-specific *Wdr72*-deficient mice, we followed the strategy described by Skarnes *et al.*^25^ Specifically, Wdr72^tm1a(KOMP)Wtsi^ mice were first crossed with *Flp recombinase* (*FlpE*) deleter mice (JAX stock #016226)^26^, which express the *FlpE* ubiquitously, to remove the FRT-flanked *lacZ* and neomycin resistance cassettes, resulting in the conditional *tm1c* (i.e., *Wdr72^fl/fl^*) allele. The following primer pairs were used to identify the *tm1c* allele or the allele with *lacZ* and neomycin resistance cassette: F1 5’-GGCGTAGGTTTTTGTGAGGATG, R1 3’-GGTGCACTTGTTTCTCATGTCC, R2 3’-GCTAAGGCAGGAGGATCACATGTTG. *Wdr72^fl/fl^* mice were subsequently bred with heterozygous *Pax8-Cre* (JAX stock #028196)^27^ knock-in mice to mediate Cre-dependent excision of exon 3, generating kidney-specific *Wdr72* knockout animals (*Wdr72^fl/fl^;Pax8-Cre^+^*), with littermate *Wdr72^fl/fl^;Pax8-Cre^-^*mice serving as controls. For all experiments, only male mice were used, and age-matched littermates served as controls. Genotyping was performed using genomic DNA extracted from ear punch biopsies, analyzed by PCR or RT-PCR with the following primer pairs: Cre-F 5’-GAACGCACTGATTTCGACCA, Cre-R 3’-AACCAGCGTTTTCGTTCTGC. To verify kidney-specific deletion of *Wdr72*, RT-PCR was performed using total RNA isolated from kidney tissue and the following pairs: Wdr72-F 5’-CACGGCCATCATGATCACAG, Wdr72-R 3’-TGCCAAACATGTCACCGA.

### Metabolic Cage Experiments and Acid-Loading Protocols

Male *Wdr72^fl/fl^;Pax8-Cre^+^* and *Wdr72^fl/fl^;Pax8-Cre^-^*littermates aged 11-17 weeks were placed individually in metabolic cages (Techniplast, Italy) for 17 hours to collect urine under mineral oil with the addition of ProClin300 (Merck, Germany) to prevent bacterial growth. Mice had free access to a standard rodent diet (wet food) and drinking water containing 2% sucrose. Metabolic cages were housed within a temperature-controlled cabinet set to 26-28 °C to minimize cold stress during the collection period. For acute acid loading, the drinking water was replaced with 0.28 M NH₄Cl supplemented with 2% sucrose for 17 hours during metabolic cage housing. For chronic acid loading, mice received 0.28 M NH₄Cl in 2% sucrose for 6 days while maintained in standard cages, followed by 17 hours in metabolic cages under the same treatment conditions. Mice were monitored daily throughout the experiment to assess general health and hydration status. At the end of each experimental period, mice were anesthetized by intraperitoneal injection of ketamine (100 mg/kg) and xylazine (20 mg/kg), and blood was collected via cardiac puncture into heparinized syringes. Plasma and urine were analyzed for clinical chemistry parameters using an AU480 Chemistry Analyzer (Beckman Coulter) and an ABL800 FLEX blood gas analyzer (Radiometer).

### Urine pH, Ammonium, and Titratable Acid Measurements

Immediately after the urine collection period in metabolic cages, urine samples were retrieved from beneath the mineral oil layer and analyzed without delay to prevent alterations due to air exposure. Urine pH was measured using a micro pH electrode. Ammonium (NH₄⁺) and titratable acids minus bicarbonate ([TA – HCO₃⁻]) were determined as previously described.^28^ Titration accuracy was verified using standard solutions of NaHCO₃ and NH₄Cl to generate calibration curves (**Supplemental Figure 1**).

Net acid excretion (NAE) was calculated according to the following formula: NAE = TA – HCO₃⁻ + NH₄⁺.

The acid–base score (AB score) was calculated as: AB score = (log₁₀[NH₄⁺] × pH³·⁶) / 40.^29^

### Tissue Collection and Fixation

Male *Wdr72^fl/fl^;Pax8-Cre^+^* mice and *Wdr72^fl/fl^;Pax8-Cre^-^* littermates (n=3) aged 8-9 weeks were anesthetized with ketamine (100 mg/kg) and xylazine (20 mg/kg) and perfused transcardially with 4% paraformaldehyde (PFA) in PBS. Procedures were conducted under license G 066/22. Kidneys were collected, post-fixed, and embedded in paraffin for immunostaining of kidney cell markers. For mice housed in metabolic cages, kidneys were collected immediately after the experiment, snap-frozen in liquid nitrogen, and stored at −80 °C for subsequent RNA and protein analysis (qPCR and Western blot).

### Immunofluorescence and confocal microscopy

Paraffin-embedded tissue blocks were sectioned at 5 µm, deparaffinized, and rehydrated. Sections were subjected to antigen retrieval in a pressure cooker using a retrieval buffer (10 mM Tris Base, 1 mM EDTA, 0.05% Tween-20, pH 9.0) for 20 minutes. Following retrieval, sections were washed in PBS containing 0.05% Tween-20 and incubated with blocking solution (PBS, 10% donkey serum, 0.3% Triton X-100) at room temperature for 1 hour.

The following primary antibodies were used: rabbit anti–Atp6v1b1 (1:1000), guineapig anti-Pendrin (1:1000), mouse anti-Atp6v1b1 B1 (68219-1-IG-20UL, proteintech, USA), goat anti-AQP2 (1:1000) (Novus Biologicals, USA), rabbit-anti-WDR72 (1:500) (ab188518, abcam, UK). The primary antibodies were incubated overnight at 4°C. Sections were subsequently washed and incubated with the appropriate secondary antibodies at room temperature for 1 hour. The nuclei were stained with DAPI. Then, the sections were mounted and imaged on a Keyence BZ-X800 microscope, a Zeiss Observer7 microscope, a Zeiss LSM 780 AxioObserver confocal microscope, or at a Leica Stellaris 8 confocal microscope. Images were then analysed using ImageJ (version Java 1.8.0_322 (64bit)).

### Quantification of the collecting duct cell composition

To assess distal tubule cell composition, kidney sections of *Wdr72^fl/fl^;Pax8 Cre^+^* and *Wdr72^fl/fl^;Pax8 Cre^-^* mice subjected to standard diet or acid load diet were analyzed by immunofluorescence staining (n = 3 per condition). For each animal, 10–12 randomly selected, non-overlapping cortical fields were imaged at 20× magnification. Cell counts were performed manually in a blinded manner. PCs were defined as aquaporin-2 (Aqp2)–positive, and ICs as Atp6v1b1–positive. IC subtypes were identified by dual labeling for Atp6v1b1 and pendrin: Type A ICs were Atp6v1b1-positive/pendrin-negative, and non-type A ICs were double-positive. Only tubules containing both Atp6v1b1 and pendrin-positive cells were included in the cell subtype analysis to ensure collecting duct selection. The investigator was blinded to genotype but not to treatment condition.

### Cell Size Analysis and Atp6v1b1 subcellular localization

To assess Atp61v1b1 subcellular localization, kidney sections from *Wdr72^fl/fl^;Pax8 Cre^+^* and *Wdr72^fl/fl^;Pax8 Cre^-^* mice were immunostained for Atp6v1b1 and pendrin, and imaged by confocal microscopy (Zeiss LSM 780 AxioObserver, Plan-Apocromat 63X/1.40 Oil DIC M27). Three mice per group were analyzed. At least 20 cortical type A ICs (Atp6v1b1-positive/pendrin-negative) and 30 cortical non-type A ICs (Atp6v1b1-positive/pendrin-positive) were evaluated per mouse. Only cells with a clearly visible nucleus were included to ensure consistent orientation. To measure cell size, the freehand selection tool in ImageJ was used to manually trace the perimeter of each IC. Area was calculated automatically by ImageJ for each cell.

### Western blot

For protein quantification, whole kidneys were homogenized and centrifuged at 1000 × g for 5 min, and the supernatant was collected. Protein lysates (100 µg) were denatured at 95 °C for 3 min before electrophoresis. Cell fractionation was performed by sequential centrifugation of whole kidney homogenate at 4°C. Samples were first spun twice at 1000 × g for 5 min, then, the resulting supernatant was centrifuged at 6000 × g for 15 min. A portion of this supernatant was kept as a total protein sample, while the remainder was ultracentrifuged at 100 000 × g for 1 h to separate cytosolic (supernatant) and membrane (pellet) fractions. Primary antibodies included rabbit anti–Atp6v1b1(1:1000), rabbit anti-Glutamine Sythetase (1:1500) (ab73593, Abcam, USA), rabbit anti-PEPCK (1:1000) (10004943, Cayman Chemical, USA) and mouse anti-β-tubulin (1:1000) (T5201-200UL, Sigma, USA); the appropriate HRP-conjugated secondary antibodies were used for detection.

### Quantitative reverse transcription PCR

Total RNA was extracted from kidney tissues using TRIzol reagent (Life Technologies, USA) and further purified with the RNeasy Mini Kit (Qiagen, Hilden, Germany). First-strand cDNA synthesis was performed using the High-Capacity cDNA Reverse Transcription Kit (Applied Biosystems, Waltham, USA). Quantitative PCR was conducted with PowerUp SYBR Green Master Mix (Applied Biosystems, Waltham, USA) in technical duplicates. Gene expression levels were normalized to *Tbp* expression, and relative expression was calculated as previously described.^30^

### Statistical analysis

Data are presented as the mean ± standard error of the mean (SEM). Comparisons between two groups were performed using unpaired two-tailed Welch’s t-test. For experiments with two independent variables (e.g., genotype and treatment), two-way analysis of variance (ANOVA) followed by appropriate post hoc tests was used. A p-value < 0.05 was considered statistically significant. All analyses were performed using GraphPad Prism version 9 or 10.

## Results

### Generation of a kidney-specific *Wdr72* knockout mouse model

Pathogenic *WDR72* variants have been implicated in dRTA and AI in humans, with 26 unrelated families harboring 21 distinct mutations reported to date (**Supplemental Figure 2**). ^10–16^ Single-cell RNA sequencing demonstrated that *Wdr72* is predominantly expressed in ICs of the CD (**Figure 1A, Supplemental Figure 3**).^22,31,32^ This cell type is critically involved in dRTA pathogenesis. Consistent with this data, immunofluorescence analysis confirmed Wdr72 protein localization in both type A ICs and non-type A ICs (**Figure 1B**).

**Figure 1.**
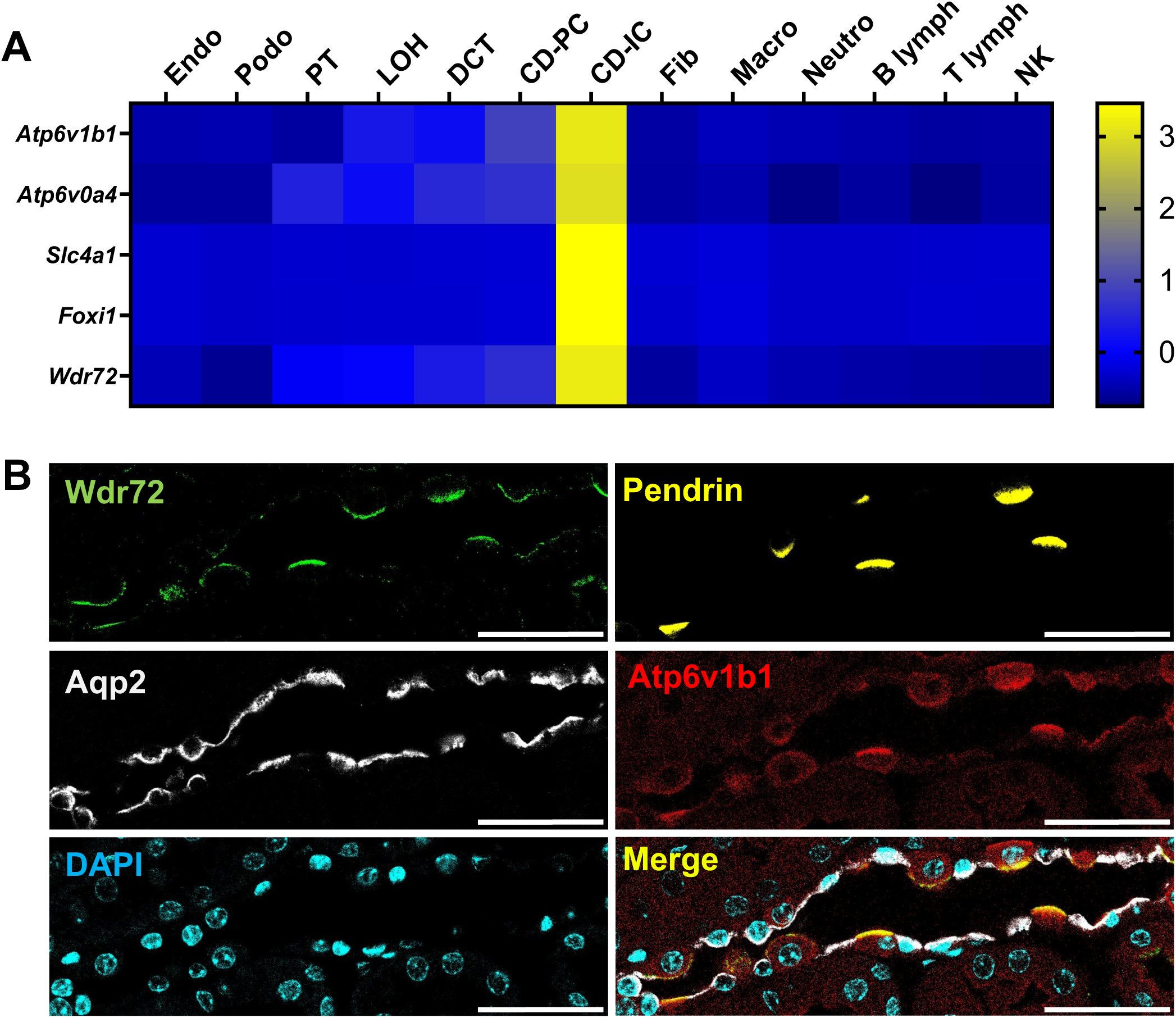
Renal expression of *Wdr72*. Cell type-specific expression of *Wdr72*. The heat map is based on *Z*-scores, modified from Park et al.^22^ Each column represents a cell type. Five dRTA genes (*Atp6v1b1*, *Atp6v0a4*, *Slc4a1*, *Foxi1, Wdr72*) are indicated. Note, dRTA genes have high expression in IC, including *Wdr72*. (**B**) Immunofluorescence images taken from cortical CD of *Wdr72^fl/fl^;Pax8 Cre^-^* stained with antibodies against Wdr72, Atp6v1b1, Aqp2, and pendrin. Wdr72 co-localizes with Atp6v1b1, and/or pendrin (type A and non-type A IC), not with the PC marker Aqp2. Nuclei are counterstained with DAPI. Endo, endothelium; Podo, podocyte; PT, proximal tubule; LOH, loop of Henle; CD-PC, collecting duct principal cell; CD-IC, collecting duct intercalating cell; Fib, fibroblast; Macro, macrophages; Neutro, neutrophil; lymph, lymphocytes; NK, natural killer cells. Scale bar 100 µm; n^IF^=3.

To investigate the role of *Wdr72* in renal acid–base homeostasis, we generated a kidney-specific *Wdr72* knockout mouse. A conditional approach was chosen to circumvent potential confounding effects associated with the global *Wdr72* knockout phenotype, including enamel defects, leading to poor feeding and reduced body weight.^33^ *Wdr72^tm1a(KOMP)Wtsi^*mice were crossed with *FlpE* deleter mice to excise the FRT-flanked *lacZ* and neomycin selection cassettes, yielding the tm1c allele (=*Wdr72^fl/fl^* **Figure 2A**).^25^ Resulting *Wdr72^fl/fl^* mice were viable and fertile and exhibited no overt anatomical or physiological abnormalities.

**Figure 2.**
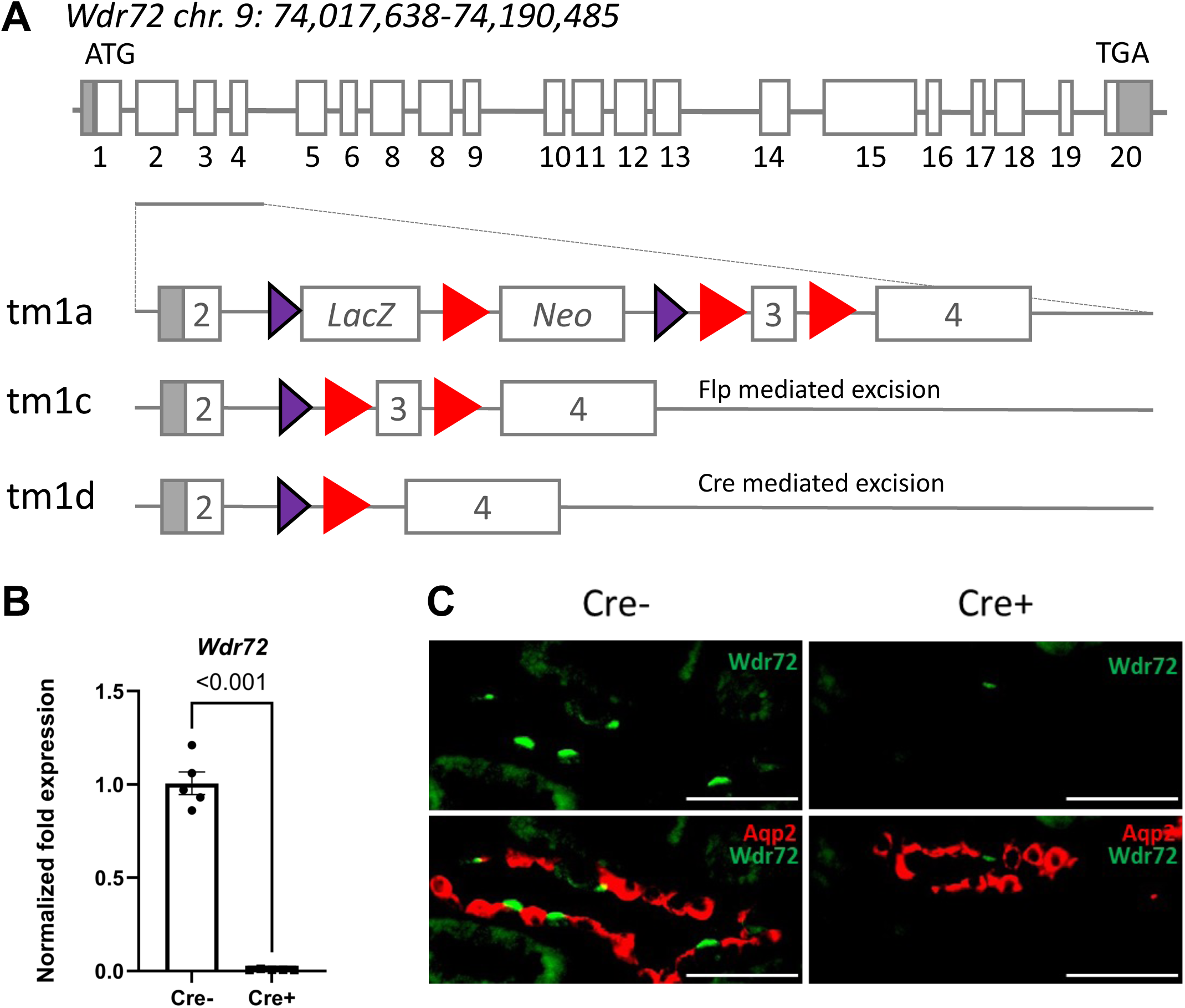
Generation of a kidney-specific *Wdr72* knockout mouse model. (**A**) Schematic representation of the murine *Wdr72* wild-type (WT) gene located on chromosome 9. White boxes indicate coding exons, and grey boxes indicate noncoding regions. The ATG start codon and TGA stop codon are shown. Below, the strategy for conditional knockout using the “knockout-first, conditional-ready” approach is illustrated.^25^ The purple triangles indicate the FRT sites, while red triangles indicate the LoxP sites. (**B**) RT-PCR analysis of kidneys from *Wdr72^fl/fl^;Pax8 Cre^-^* vs. *Wdr72^fl/fl^;Pax8 Cre^+^* mice showing loss of *Wdr72* expression in knockout samples. (**C**) Representative immunofluorescence images of cortical CD from *Wdr72^fl/fl^;Pax8 Cre^-^* vs. *Wdr72^fl/fl^;Pax8 Cre^+^* mice stained with an anti-Wdr72 antibody as well as anti-Aqp2. The *Wdr72^fl/fl^;Pax8 Cre^+^* mice exhibit reduced Wdr72 signal in IC compared with controls. Data were analyzed using unpaired *t*-test; values are reported as means +/- SEM; p-values <0.05 are considered significant; n^qPCR^=5. Scale bar 50 µm; n^IF^=3.

Next, *Wdr72^fl/fl^* were crossed with *Pax8-Cre* transgenic mice to achieve kidney specific deletion of exon 3 flanked by loxP sites, resulting in kidney-specific *Wdr72* knockout animals (tm1d, *Wdr72^fl/fl^*;*Pax8 Cre^+^*). *Wdr72^fl/fl^;Pax8 Cre^+^* mice were also viable and indistinguishable from Cre-negative littermates in body weight and general appearance under standard housing and diet conditions (**Supplemental Table 1**). Offspring were born at expected Mendelian ratios. PCR genotyping confirmed successful recombination of the *Wdr72* locus. Quantitative RT-PCR demonstrated the absence of full-length *Wdr72* transcripts in kidneys from *Wdr72^fl/fl^;Pax8 Cre^+^* mice (**Figure 2B**). Consistently, immunofluorescence microscopy revealed markedly reduced Wdr72 protein in kidneys from *Wdr72^fl/fl^;Pax8 Cre^+^* animals (**Figure 2C**), whereas a robust signal was detected in *Wdr72^fl/fl^;Pax8 Cre^-^* controls within ICs (**Figure 1B, 2C**).

### Kidney-specific knockout of *Wdr72* results in acid-base imbalance

To assess the impact of renal *Wdr72* loss on systemic acid–base balance, we analyzed blood and urine parameters in male *Wdr72^fl/fl^;Pax8 Cre^+^* and *Wdr72^fl/fl^;Pax8 Cre^-^* mice under baseline conditions with free access to a standard rodent diet and water. Body weight, food and water intake, as well as urine output were comparable between genotypes (**Supplemental Table 1**). However, *Wdr72^fl/fl^;Pax8 Cre^+^* had significantly higher urine pH levels compared to *Wdr72^fl/fl^;Pax8 Cre^-^* controls. This was accompanied by a reduction in net acid excretion (NAE) and titratable acids (**Figure 3A-C**, **Table 1**), indicating a potential urinary proton loss in the knockout mice. Interestingly the urinary NH₄⁺ excretion remained unchanged under standard diet (**Figure 3D**), while the AB-Score was increased (**Table 1**), indicating an altered relation between urine pH and NH₄⁺ excretion.^29^ No differences in blood pH, bicarbonate (HCO₃⁻), base excess (BE) or pCO₂ levels were detected between *Wdr72^fl/fl^;Pax8 Cre^+^* and control mice under standard diet conditions (**Table 2**). In addition, renal function parameters (serum creatinine and urea) as well as serum and urinary electrolyte levels were not altered between the two groups (**Table 2**). Thus, *Wdr72^fl/fl^;Pax8 Cre^+^* mice show an alkalotic urinary phenotype consistent with incomplete dRTA, as reported for other murine models harboring mutations in human dRTA genes.^34,35^

**Figure 3.**
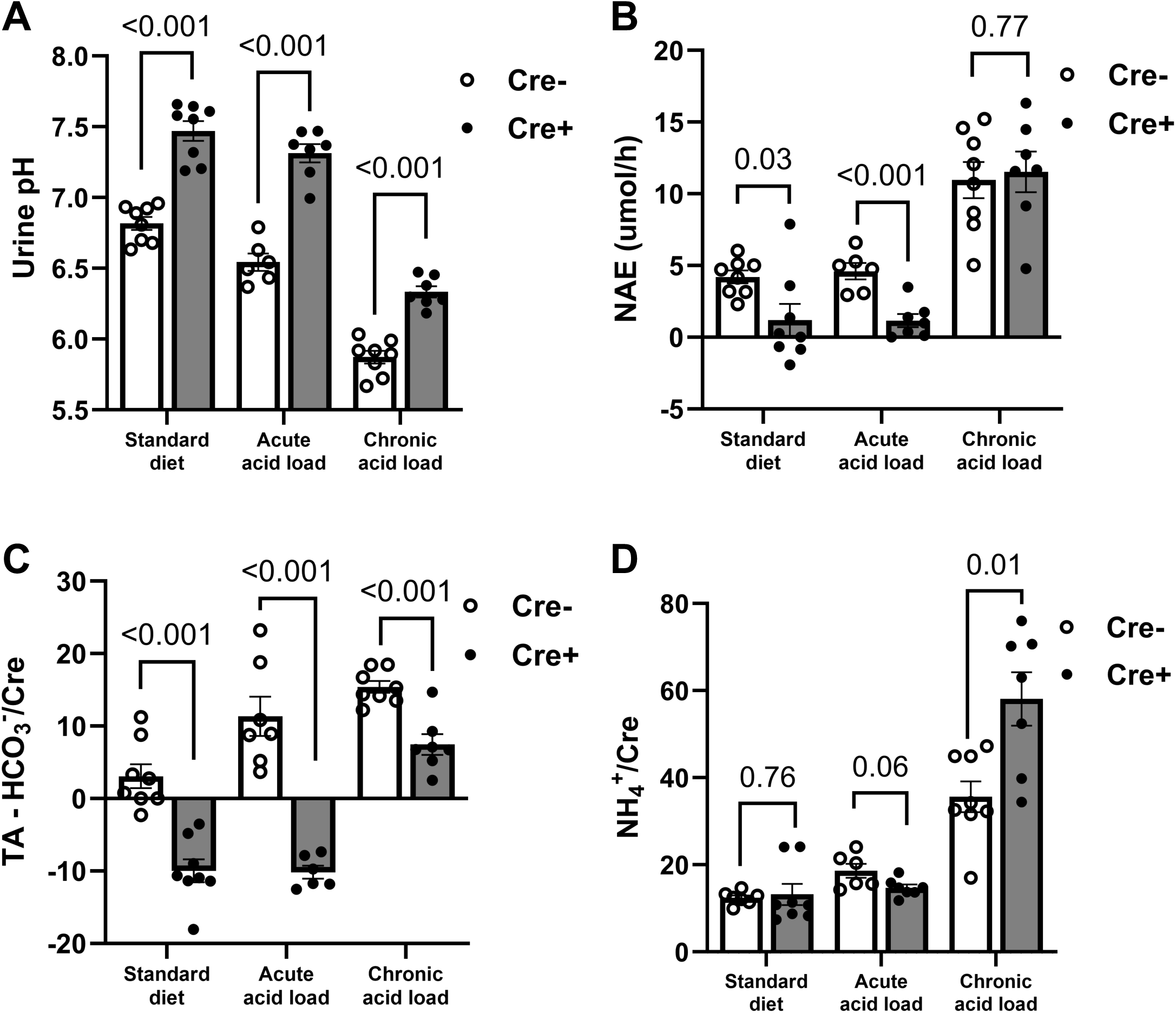
Kidney-specific knockout of *Wdr72* results in acid-base imbalance. *Wdr72^fl/fl^;Pax8 Cre^+^* mice show reduced urinary acid secretion compared with control (*Wdr72^fl/fl^;Pax8 Cre^-^*) mice under standard diet, acute acid load, and chronic acid load conditions. This reduction is reflected by higher urinary pH (**A**), lower net acid excretion (NAE) (A), and decreased titratable acidity (TA) (**C**). (**D**) Following chronic acid load, *Wdr72^fl/fl^;Pax8 Cre*^+^ mice exhibit increased urinary NH₄⁺ excretion compared with controls, resulting in compensation of NAE despite reduced TA. Data were analyzed using unpaired *t*-test; values are reported as means +/- SEM; p-values <0.05 are considered significant; n=6-8.

**Table 1.** Summary of urine data collected under standard diet, acute and chronic acid load condition. Urine was collected from *Wdr72^fl/fl^;Pax8 Cre^+^* (Cre+) vs. *Wdr72^fl/fl^;Pax8 Cre^-^* (Cre-) mice under the indicated conditions. Statistical analyses were performed using unpaired t-tests; values are reported as means +/- SEM; p-values <0.05 were considered significant; n=6-8. FE, fractional excretion; NAE, net acid excretion; TA, titrable acid; CRE, creatinine.

| Urine parameters<br>(unit) | Standard diet |  |  |  |  | Acute acid load |  |  |  |  | Chronic acid load |  |  |  |  |
| --- | --- | --- | --- | --- | --- | --- | --- | --- | --- | --- | --- | --- | --- | --- | --- |
|  | Cre- |  | Cre+ |  | p-Value | Cre- |  | Cre+ |  | p-Value | Cre- |  | Cre+ |  | p-Value |
|  | mean | ± SEM | mean | ± SEM |  | mean | ± SEM | mean | ± SEM |  | mean | ± SEM | mean | ± SEM |  |
| FE UREA (%) | 20.88 | 2.56 | 22.86 | 2.19 | 0.59 | 29.10 | 2.56 | 25.78 | 1.55 | 0.33 | 31.69 | 2.82 | 31.38 | 3.79 | 0.95 |
| FE Ca<sup>+2</sup> (%) | 0.30 | 0.05 | 0.14 | 0.01 | 0.02 | 0.29 | 0.02 | 0.15 | 0.01 | <0.001 | 0.37 | 0.01 | 0.29 | 0.05 | 0.22 |
| FE PHOS (%) | 5.36 | 0.69 | 3.37 | 0.62 | 0.07 | 7.06 | 0.19 | 4.13 | 0.56 | 0.004 | 5.91 | 0.46 | 3.44 | 0.43 | 0.004 |
| FE Mg<sup>+2</sup> (%) | 6.83 | 1.17 | 5.62 | 0.50 | 0.40 | 9.02 | 0.65 | 4.01 | 0.62 | <0.001 | 10.13 | 0.86 | 6.04 | 0.43 | 0.004 |
| FE Na<sup>+</sup> (%) | 0.33 | 0.04 | 0.35 | 0.03 | 0.72 | 0.42 | 0.02 | 0.40 | 0.02 | 0.69 | 0.25 | 0.02 | 0.24 | 0.04 | 0.79 |
| FE K<sup>+</sup> (%) | 14.57 | 0.74 | 16.07 | 1.02 | 0.30 | 20.31 | 0.92 | 22.14 | 1.44 | 0.35 | 12.55 | 1.30 | 14.37 | 1.69 | 0.45 |
| FE Cl<sup>-</sup> (%) | 41.41 | 1.65 | 40.18 | 1.96 | 0.67 | 59.98 | 3.06 | 55.49 | 1.76 | 0.28 | 67.91 | 7.08 | 84.74 | 8.97 | 0.21 |
| Urine pH | 6.82 | 0.04 | 7.47 | 0.07 | <0.001 | 6.54 | 0.06 | 7.31 | 0.06 | <0.001 | 5.87 | 0.04 | 6.33 | 0.04 | <0.001 |
| AB score | 23.31 | 1.79 | 34.55 | 1.66 | 0.002 | 28.92 | 0.90 | 41.13 | 0.83 | <0.001 | 26.29 | 0.56 | 38.40 | 0.64 | <0.001 |
| NAE (umol/h) | 4.21 | 0.41 | 0.03 | 1.05 | 0.03 | 4.60 | 0.52 | 1.16 | 0.43 | <0.001 | 10.96 | 1.18 | 11.53 | 1.31 | 0.77 |
| [TA - HCO<sub>3</sub>]/CRE | 3.08 | 1.54 | -9.97 | 1.48 | <0.001 | 11.34 | 2.50 | -10.17 | 0.82 | <0.001 | 15.41 | 0.75 | 7.47 | 1.33 | <0.001 |
| NH<sub>4</sub><sup>+</sup>/CRE | 12.42 | 0.52 | 13.22 | 2.27 | 0.76 | 18.61 | 1.47 | 14.72 | 0.70 | 0.06 | 35.64 | 3.29 | 58.09 | 5.67 | 0.01 |

**Table 2.** Summary of blood gas analysis collected under standard diet, acute and chronic acid load condition. Data was collected from *Wdr72^fl/fl^;Pax8 Cre^+^* (Cre+) vs. *Wdr72^fl/fl^;Pax8 Cre^-^* (Cre-) mice under the indicated conditions. Statistical analyses were performed using unpaired t-tests; values are reported as means +/- SEM; p-values <0.05 are considered significant; n=6-7. *C*, calculated; ABE, actual base excess; SBE, standard base excess; Hct, hematocrit; CRE, creatinine.

| Blood parameters<br>(unit) | Standard diet |  |  |  |  | Acute acid load |  |  |  |  | Chronic acid load |  |  |  |  |
| --- | --- | --- | --- | --- | --- | --- | --- | --- | --- | --- | --- | --- | --- | --- | --- |
|  | Cre- |  | Cre+ |  | p-Value | Cre- |  | Cre+ |  | p-Value | Cre- |  | Cre+ |  | p-Value |
|  | mean | ± SEM | mean | ± SEM |  | mean | ± SEM | mean | ± SEM |  | mean | ± SEM | mean | ± SEM |  |
| pH | 7.20 | 0.01 | 7.16 | 0.02 | 0.12 | 7.10 | 0.03 | 7.08 | 0.01 | 0.48 | 7.20 | 0.02 | 7.15 | 0.01 | 0.048 |
| pCO<sub>2</sub> (mmHg) | 69.19 | 1.75 | 67.87 | 2.10 | 0.66 | 63.46 | 1.43 | 62.36 | 2.25 | 0.71 | 67.36 | 1.60 | 74.00 | 1.48 | 0.02 |
| cHCO<sub>3</sub><sup>-</sup>(P)<sub>c</sub> (mM) | 25.53 | 1.04 | 23.50 | 0.93 | 0.20 | 19.11 | 1.21 | 17.63 | 0.83 | 0.37 | 25.40 | 1.18 | 24.53 | 0.56 | 0.55 |
| cHCO<sub>3</sub><sup>-</sup>(Pst)<sub>c</sub> (mM) | 19.43 | 0.83 | 17.70 | 0.77 | 0.18 | 14.31 | 1.09 | 13.09 | 0.54 | 0.38 | 19.51 | 0.94 | 18.07 | 0.53 | 0.25 |
| ABEc (mM) | -4.11 | 1.13 | -6.43 | 1.09 | 0.20 | -11.70 | 1.70 | -13.43 | 0.89 | 0.43 | -3.97 | 1.29 | -5.90 | 0.67 | 0.25 |
| SBEc (mM) | -1.73 | 1.11 | -4.03 | 1.07 | 0.19 | -9.07 | 1.51 | -10.83 | 0.88 | 0.37 | -1.73 | 1.33 | -3.32 | 0.68 | 0.35 |
| Hctc (%) | 44.44 | 0.41 | 43.95 | 0.61 | 0.55 | 44.30 | 0.61 | 43.96 | 0.41 | 0.68 | 43.20 | 0.81 | 41.78 | 1.06 | 0.35 |
| UREA (mM) | 11.04 | 0.62 | 10.68 | 0.50 | 0.68 | 11.56 | 0.80 | 12.77 | 0.66 | 0.30 | 8.86 | 0.59 | 10.72 | 0.86 | 0.13 |
| Ca<sup>2+</sup> (mM) | 2.40 | 0.06 | 2.51 | 0.07 | 0.28 | 2.56 | 0.05 | 2.49 | 0.03 | 0.31 | 2.55 | 0.02 | 2.38 | 0.02 | <0.001 |
| GLUC (mM) | 22.25 | 1.48 | 21.23 | 1.66 | 0.68 | 16.76 | 1.35 | 16.66 | 0.87 | 0.96 | 20.64 | 0.99 | 16.67 | 1.10 | 0.03 |
| PHOS (mM) | 2.74 | 0.08 | 3.17 | 0.20 | 0.10 | 2.80 | 0.09 | 2.96 | 0.03 | 0.15 | 2.97 | 0.08 | 3.58 | 0.22 | 0.05 |
| Mg<sup>+2</sup> (mM) | 1.20 | 0.03 | 1.20 | 0.03 | 0.93 | 1.21 | 0.05 | 1.18 | 0.03 | 0.67 | 0.99 | 0.03 | 1.06 | 0.03 | 0.15 |
| Na<sup>+</sup> (mM) | 143.63 | 1.13 | 143.50 | 0.81 | 0.93 | 148.57 | 1.51 | 149.14 | 1.39 | 0.80 | 145.86 | 0.55 | 149.33 | 0.65 | 0.004 |
| K<sup>+</sup> (mM) | 4.74 | 0.20 | 4.25 | 0.11 | 0.07 | 4.37 | 0.13 | 4.10 | 0.18 | 0.27 | 4.99 | 0.15 | 4.23 | 0.16 | 0.01 |
| Cl<sup>-</sup> (mM) | 112.00 | 0.85 | 111.67 | 0.56 | 0.77 | 120.57 | 1.40 | 121.43 | 1.65 | 0.72 | 112.71 | 0.83 | 115.50 | 0.74 | 0.04 |
| CRE (mM) | 0.0085 | 0.0008 | 0.010 | 0.0005 | 0.19 | 0.010 | 0.0005 | 0.011 | 0.0003 | 0.47 | 0.0096 | 0.0002 | 0.010 | 0.0004 | 0.19 |

To evaluate the capacity of *Wdr72^fl/fl^;Pax8 Cre^+^*mice to respond to acid challenge, we administered 0.28 M ammonium chloride (NH₄Cl) in the drinking water. We tested both an acute (17 h) and a chronic (7 days) acid challenge. No differences were observed between genotypes in body weight, urine volume, or food and water intake under acute and chronic acidified conditions (**Supplemental Table 1**). As expected, NH₄Cl ingestion increased urinary acidity in both *Wdr72^fl/fl^;Pax8 Cre^-^* controls and *Wdr72^fl/fl^;Pax8 Cre^+^* mice (**Figure 3A**, **Table 2**). However, urinary pH remained significantly higher in *Wdr72^fl/fl^;Pax8 Cre^+^* mice compared to controls under both acute and chronic acid loading (**Figure 3A**, **Table 2**). This indicates a defect in the maximal urinary acidification in response to acid challenge, a hallmark of incomplete distal renal tubular acidosis. Titratable acids were significantly reduced in acute and chronic acid load in the knockout mice, again emphasizing a potential proton secreting defect (**Figure 3C**). Interestingly, the *Wdr72^fl/fl^;Pax8 Cre^+^* mice had an increased urinary NH₄⁺ excretion under chronic acid load accompanied by a rise in NAE (**Figure 3B, D**). This may suggest enhanced ammoniagenesis as a compensatory response to impaired proton secretion.

The acute acid load induced mild systemic acidosis in the *Wdr72^fl/fl^;Pax8 Cre^-^* controls (**Table 1, Supplemental Table 2**). However, no differences in blood gas analysis were observed between the *Wdr72^fl/fl^;Pax8 Cre^+^* vs. the *Wdr72^fl/fl^;Pax8 Cre^-^* controls under acute acid load (**Table 2**). The urinary excretion of calcium, phosphate, and magnesium was reduced in *Wdr72^fl/fl^;Pax8 Cre^+^* mice compared to controls (**Table 2**).

Following 7 days of chronic acid loading, *Wdr72^fl/fl^;Pax8 Cre^+^* exhibited a modest but significant decrease in systemic blood pH compared to *Wdr72^fl/fl^;Pax8 Cre^-^* controls (**Table 2**). This, however, was accompanied by a slightly increased pCO_2_, but also increased serum sodium, chloride and phosphate levels as well as decreased serum potassium and calcium levels, without a change of the hematocrit (**Table 2**). Additionally, the fractional excretion of phosphate and magnesium were reduced in *Wdr72^fl/fl^;Pax8 Cre^+^* vs. *Wdr72^fl/fl^;Pax8 Cre^-^* mice under chronic acid load (**Table 1**).

As nephrocalcinosis is observed in approximately 77% of patients with *WDR72* mutations^10^, we performed Alizarin Red and von Kossa staining on kidney sections from 9- and 23- week-old mice. However, no nephrocalcinosis was detected in either *Wdr72^fl/fl^;Pax8 Cre^+^*nor *Wdr72^fl/fl^;Pax8 Cre^-^* control mice (**Supplemental Figure 3**).

Overall, our data indicate that *Wdr72^fl/fl^;Pax8 Cre^+^* mice fail to adequately respond to an oral acid challenge, highlighting a potential role for Wdr72 in incomplete dRTA, while not fully recapitulating the human *WDR72* phenotype.

### Loss of *Wdr72* increases ammoniagenesis markers in kidneys

As chronic acid load resulted in increased urinary NH₄⁺ excretion and elevated NAE, despite persistently reduced TA, we next assessed markers of renal ammoniagenesis by Western blot. Interestingly, Phosphoenolpyruvate carboxykinase (Pepck) protein levels were increased under standard diet and chronic acid load in the kidneys of *Wdr72^fl/fl^;Pax8 Cre^+^* mice compared to *Wdr72^fl/fl^;Pax8 Cre^-^* controls, while Glutamine synthetase (GS) protein levels were decreased in the kidneys of *Wdr72^fl/fl^;Pax8 Cre^+^* mice compared to *Wdr72^fl/fl^;Pax8 Cre^-^* controls (**Figure 4A-D**). This indicates an adaptive compensatory response in the *Wdr72* deficient mice to enhance glutamine catabolism and ammoniagenesis to support NH₄⁺-dependent acid excretion in the knockout mice.

**Figure 4.**
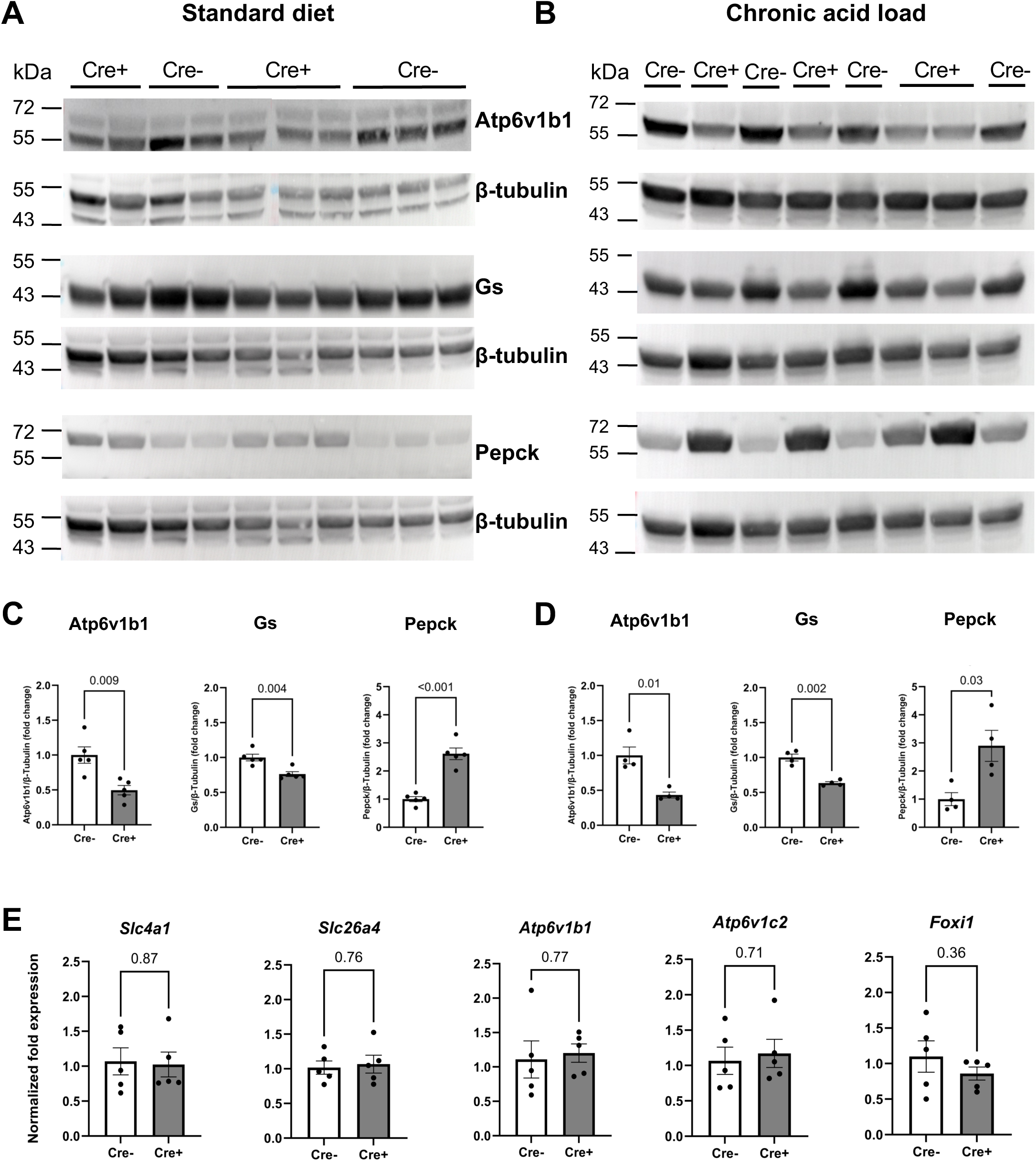
Kidney-specific loss of *Wdr72* leads to reduced Atp6v1b1 but enhanced ammoniagenesis pathways within the kidney. Western blot analysis of whole-kidney lysates from *Wdr72^fl/fl^;Pax8 Cre^-^* (Cre-) vs. *Wdr72^fl/fl^;Pax8 Cre^+^* (Cre+) mice under standard diet (**A**) and following chronic acid challenge (**B**) demonstrates reduced Atp6v1b1 protein abundance in *Wdr72^fl/fl^;Pax8 Cre^+^* kidneys, accompanied by markedly reduced protein abundance of Glutamine synthetase (Gs) and increased Phosphoenolpyruvate carboxykinase (Pepck), indicating compensatory upregulation of ammoniagenesis pathways and a potential defect in post-transcriptional trafficking or stabilization of Atp6v1b1. (**C, D**) Quantification of protein abundance relative to β-tubulin was performed using ImageJ. (**E**) qPCR analysis of *Slc4a1*, *Slc26a4*, *Atp6v1b1*, *Atp6v1c2*, and *Foxi1* mRNA expression in kidneys from *Wdr72^fl/fl^;Pax8 Cre^-^* (Cre-) vs. *Wdr72^fl/fl^;Pax8 Cre^+^* (Cre+) mice under basal conditions shows no significant differences in transcript levels. Data were analyzed with unpaired *t*-test; values are reported as means +/- SEM; p-values <0.05 are considered significant; n^WB^=4-5; n^qPCR^=5.

### Kidney-specific loss of *Wdr72* leads to impaired regulation of V-ATPase abundance or stability

To investigate potential compensatory mechanisms in *Wdr72^fl/fl^;Pax8 Cre^+^* mice, we next analyzed the expression of genes encoding several acid–base proteins within cells of the collecting duct (CD) by qPCR and Western blot. No differences were observed at the mRNA level for *Ae1*, *Foxi1*, *Atp6v1b1*, *Pendrin*, *Atp6v1c2* (**Figure 4E**). This was further supported by bulk RNA sequencing of whole-kidney RNA from *Wdr72^fl/fl^;Pax8 Cre^+^* vs. *Wdr72^fl/fl^;Pax8 Cre^-^* mice, which revealed no significant changes in the expression of acid–base–related genes (**Supplemental Figure 5**). However, Western Blot analysis of the B1 subunit of the V-ATPase, encoded by *Atp6v1b1*, a major gene implicated in monogenic dRTA, revealed a significant reduction in protein abundance in *Wdr72^fl/fl^;Pax8 Cre^+^* vs. *Wdr72^fl/fl^;Pax8 Cre^-^* mice under standard and chronic acid load conditions (**Figure 4A-D**). The V-ATPase B1 subunit is critical for proton secretion in ICs and is a known dRTA gene in mice and humans.^6,34,35^ This discrepancy between *Atp6v1b1* mRNA and protein abundance in *Wdr72^fl/fl^;Pax8 Cre^+^* mice suggests that Wdr72 regulates Atp6v1b1 post-transcriptionally, possibly by affecting protein trafficking, stability, or V-ATPase complex assembly.

### Loss of *Wdr72* in the kidney leads to an altered IC pattern

Given the reduced Atp6v1b1 protein abundance and the elevated urinary pH in *Wdr72^fl/fl^;Pax8 Cre^+^* mice, we next examined whether *Wdr72* deficiency alters the overall cellular composition of the CD (**Figure 5A-C**). We quantified the proportion of principal cells (PCs; Aqp2⁺/Atp6v1b1⁻) relative to all IC (Aqp2⁻/Atp6v1b1⁺) in the cortical CD (CCD) and inner medullary CD (IMCD) under standard diet (**Figure 5A-C**) and in the CCD under chronic acid-loading (**Supplemental Figure 6**) conditions using immunofluorescence imaging. No significant differences in the PC-to-IC ratio were observed between *Wdr72^fl/fl^;Pax8 Cre^+^* and *Wdr72^fl/fl^;Pax8 Cre^-^* under either diet (**Figure 5A-C, Supplemental Figure 6**).

**Figure 5.**
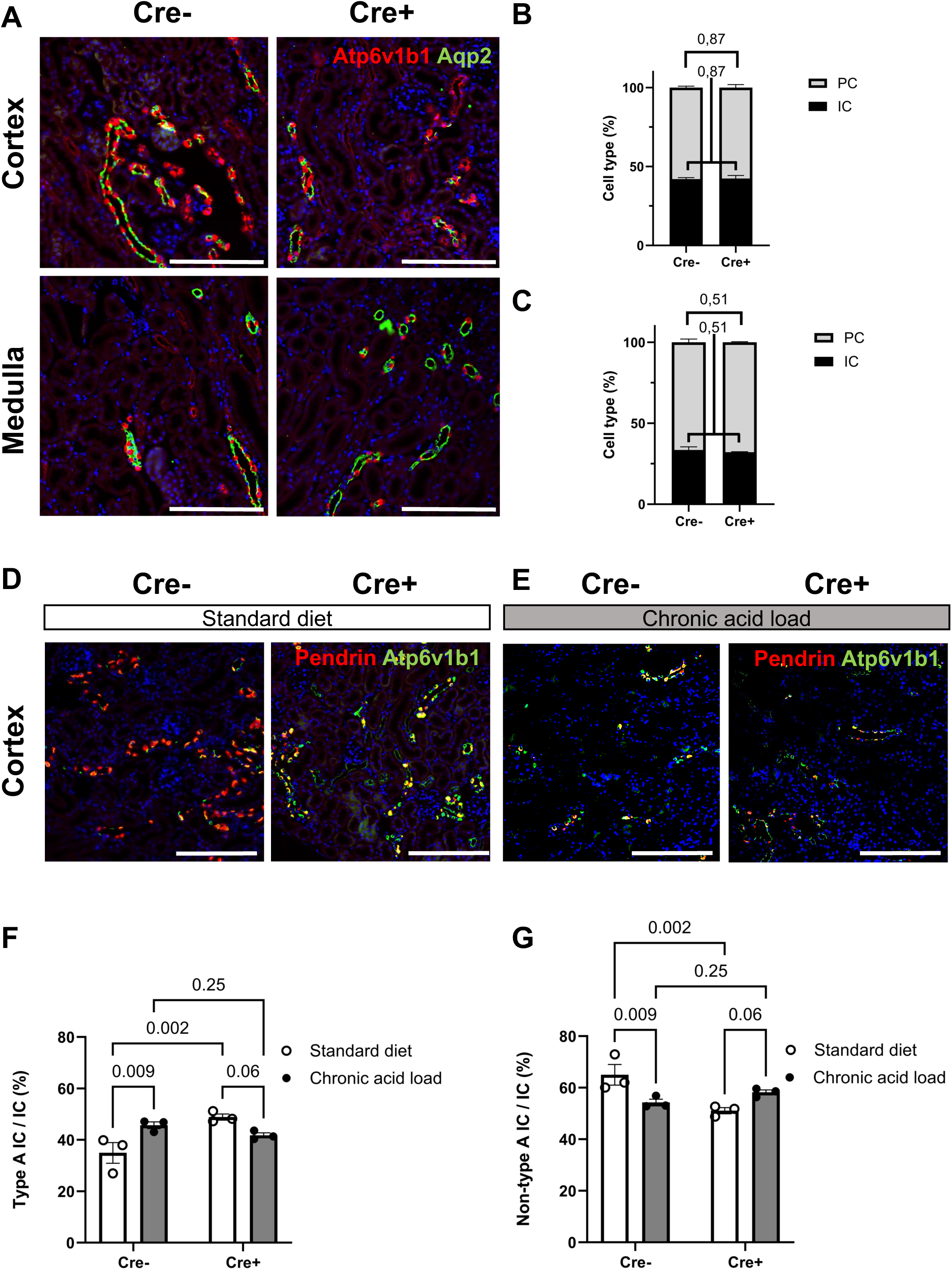
Wdr72 ablation causes altered cellular consistency in the collecting ducts. (A) Representative immunofluorescence images of cortical and medullary collecting duct cells under standard diet conditions from *Wdr72^fl/fl^;Pax8 Cre^-^* (Cre-) vs. *Wdr72^fl/fl^;Pax8 Cre^+^* (Cre+) mice. Kidneys were stained for Aqp2 (green) and Atp6v1b1 (red); nuclei were counterstained with DAPI (blue). Aqp2-positive cells were identified as PCs, and Atp6v1b1-positive cells as ICs. (**B, C)** Quantification of the IC-to-PC ratio in cortical (**B**) or inner medullary (**C**) collecting ducts shows no difference between the genotypes. (**D, E**) Representative immunofluorescence images of kidneys stained for pendrin (red) and Atp6v1b1(green) under standard diet (**D**) and chronic acid-loading (**E**) conditions from *Wdr72^fl/fl^;Pax8 Cre^-^* (Cre-) vs. *Wdr72^fl/fl^;Pax8 Cre^+^* (Cre+) mice. **(F, G)** Quantification of type A and non-type A ICs expressed as a percentage of total ICs. Note: Under basal conditions, cortical type A IC percentages were significantly higher in *Wdr72^fl/fl^;Pax8 Cre^+^* mice compared to controls. Differences between genotypes and conditions were analyzed using a two-way ANOVA test; values are reported as means +/- SEM; p-values <0.05 are considered significant; n=3. Scale bars 200 μm.

Different IC subtypes exist, with type A ICs being the primary mediators of acid secretion due to their apical localization of Atp6v1b1. To distinguish IC subtypes, we employed anti-pendrin and anti-Atp6v1b1 staining. Type A ICs are pendrin-negative, Atp6v1b1–positive cells, while non-type A ICs are pendrin-positive, Atp6v1b1–positive cells. We then analyzed these subpopulations in the kidneys of *Wdr72^fl/fl^;Pax8 Cre^+^* and *Wdr72^fl/fl^;Pax8 Cre^-^* mice under standard diet or chronic acid loading condition **(Figure 5D, E**). In control mice (*Wdr72^fl/fl^;Pax8 Cre^-^*), chronic acid loading induced an increase in the proportion of type A ICs relative to all ICs within the CCD, which was accompanied by a reduction in the non-type A IC compartment (**Figure 5F, G**). Under standard diet, *Wdr72^fl/fl^;Pax8 Cre^+^* mice exhibited a significant increase in the proportion of type A ICs among all ICs compared with control mice (**Figure 5F)**. Interestingly, following chronic acid loading, the number of type A ICs did not further increase in the knockout cohort (**Figure 5F**). Accordingly, the non-type A IC revealed a significant reduction upon acid load in the control mice, while *Wdr72^fl/fl^;Pax8 Cre^+^* mice exhibited a reduced number of non-A IC under standard diet, with no change upon acid load (**Figure 5G**). These data suggest that *Wdr72^fl/fl^;Pax8 Cre^+^* mice have an altered adaptive response of ICs under standard diet but also to chronic acid loading, potentially limiting their ability to further increase acid-secreting capacity.

### Loss of *Wdr72* in the kidney leads to changes in IC size

Altered IC morphology has been reported in other models of dRTA.^36,37^ To evaluate cellular morphology in *Wdr72^fl/fl^;Pax8 Cre^+^* mice, we measured the size of ICs in the murine CCD using immunofluorescence and confocal microscopy. We observed that non-type A ICs (**Figure 6A)** were significant smaller in *Wdr72^fl/fl^;Pax8 Cre^+^* mice compared with *Wdr72^fl/fl^;Pax8 Cre^-^* controls, whereas there was only a tendency toward smaller type A ICs in *Wdr72^fl/fl^;Pax8 Cre^+^* mice The difference was maintained upon chronic acid load, with *Wdr72^fl/fl^;Pax8 Cre^+^* kidney continuing to show smaller ICs than the controls (**Figure 6B**). The reduced size of ICs in *Wdr72^fl/fl^;Pax8 Cre^+^* together with their overall increased number, indicates a reduced cellular plasticity in response to acid challenge within the CCD.

**Figure 6.**
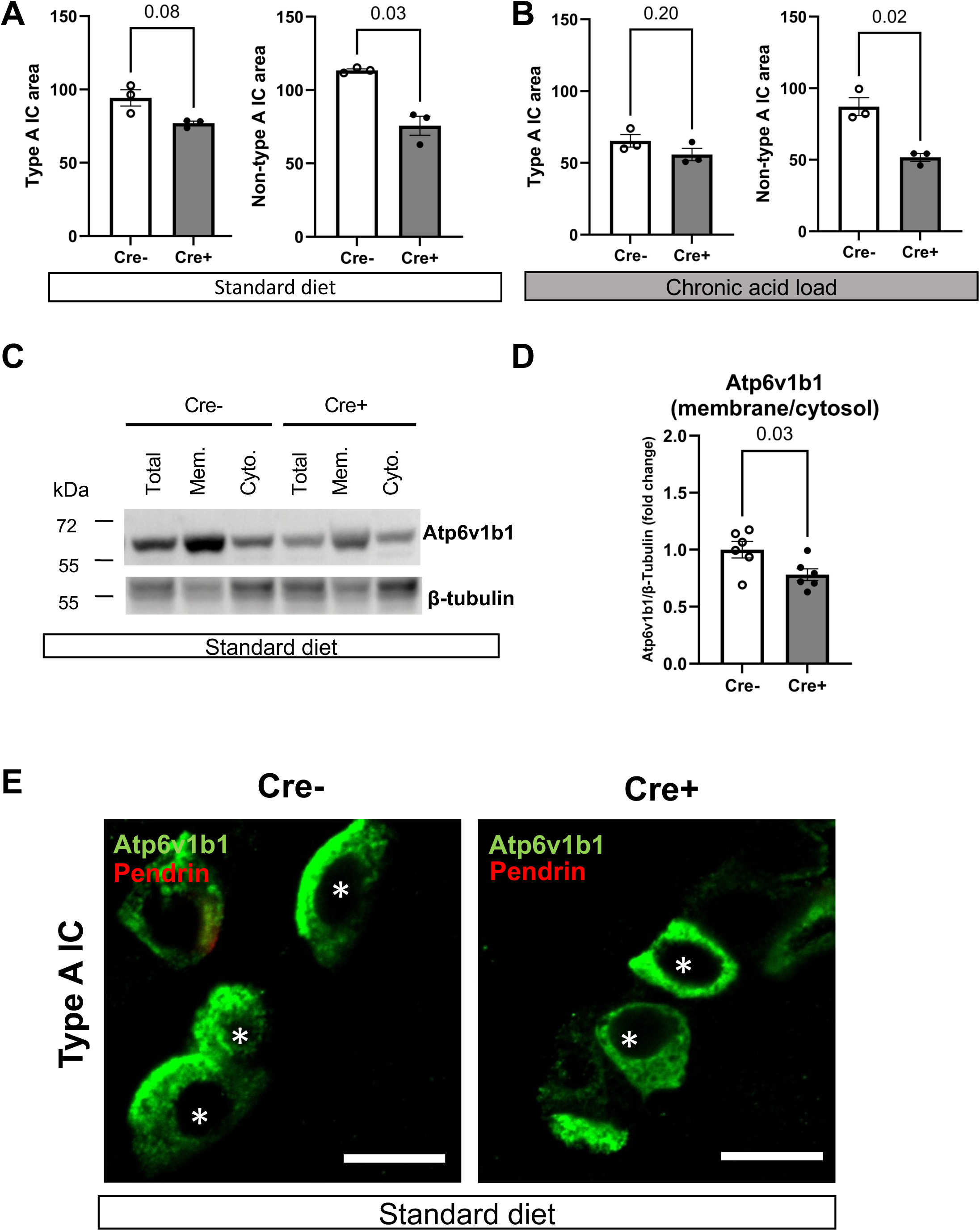
Altered Atp6v1b1 localization and cell size of intercalated cells in *Wdr72^fl/fl^;Pax8 Cre^+^* kidneys. The cell sizes of type A and non-type A IC in the cortical collecting ducts were determined under standard diet (**A**) and chronic acid load (**B**) from *Wdr72^fl/fl^;Pax8 Cre^-^* (Cre-) vs. *Wdr72^fl/fl^;Pax8 Cre^+^* (Cre+) mice. The quantification shows a reduction in the sizes of type B IC cell types with a trend in A ICs under both dietary conditions in *Wdr72^fl/fl^;Pax8 Cre^+^* (Cre+) mice. (**C**) Western blot of subcellular fractionation of *Wdr72^fl/fl^;Pax8 Cre^+^* (Cre+) vs. *Wdr72^fl/fl^;Pax8 Cre^-^* (Cre-) kidneys of mice subjected to standard diet, showing decreased membranous (mem.), cytosolic (cyto.), and total protein abundance of Atp6v1b1. (**D**) Quantification of the membrane/cytosol ratio after normalizing to beta-tubulin expression, showing a reduction in the *Wdr72^fl/fl^;Pax8 Cre^+^.* (**E**) Representative immunofluorescence images show cortical collecting ducts with type A IC from *Wdr72^fl/fl^;Pax8 Cre^-^*(Cre-) and *Wdr72^fl/fl^;Pax8 Cre^+^* (Cre+)with more cytosolic Atp6v1b1 localization.. Statistical analyses were performed using unpaired *t*-tests; values are reported as means +/- SEM; p-values <0.05 are considered significant; n=3. Scale bar 10 μm.

### Loss of *Wdr72* in the kidney prevents membranous localization of Atp6v1b1

Type A ICs can increase the abundance of V-ATPases in their apical plasma membranes. This likely occurs through trafficking of subapical V-ATPase–containing vesicles to the apical surface, or enhanced assembly of V_1_ and V_0_ subunits into functional holoenzymes, thereby augmenting proton secretory capacity. To determine whether Wdr72 affects the membranous localization of Atp6v1b1, we performed a Western blot with cytosolic and membrane protein fractions from whole kidney lysates and probed for Atp6v1b1 within the fractions (**Figure 6C-D, Supplemental Figure 7**). In the *Wdr72^fl/fl^*;Pax8 Cre^+^ kidneys, membrane, but also total and cytosolic associated Atp6v1b1 abundance was markedly reduced compared with controls, with a significantly decreased membrane/cytosol ratio compared to control under standard diet (**Figure 6C-D, Supplemental Figure 7**). Additionally, we performed a co-staining with Atp6v1b1 subunit and pendrin to identify type A ICs (pendrin^-^, Atp6v1b1^+^) in the kidneys of *Wdr72^fl/fl^;Pax8 Cre^+^*and *Wdr72^fl/fl^;Pax8 Cre^-^* mice (**Figure 6E**,). The *Wdr72^fl/fl^;Pax8 Cre^+^* displayed a more cytosolic localization of the Atp6v1b1 subunit within the type A ICs in the kidney. These data suggest that Wdr72 is essential for the stability, but also assembly or transport of the V-ATPase holoenzyme, indicated by reduced membranous Atp6v1b1 subunit expression.

## Discussion

*WDR72* pathogenic variants have been identified in families with dRTA and AI. Here, we investigated the function of Wdr72 in renal acid-base homeostasis using a kidney-specific *Wdr72* knockout mouse model. These mice exhibited features of incomplete dRTA, including elevated urinary pH, reduced titratable acid and net acid excretion, accompanied by impaired maximal urinary acidification after acid load. Within the CD, type A ICs were increased in number but reduced in size. They failed to expand upon chronic acid challenge, indicating compromised adaptive plasticity. At the protein level, the abundance and membranous localization of the B1 subunit of the V-ATPase within ICs were significantly reduced despite unaltered *Atp6v1b1* transcript levels, suggesting a defect in protein trafficking, assembly, or stabilization. Chronic acid loading induced compensatory upregulation of ammoniagenesis and NH_4_^+^ excretion, potentially reflecting an adaptive response to maintain systemic acid-base equilibrium despite impaired distal proton secretion.

Global *Wdr72* knockout mice are viable, but develop AI and marked postnatal growth retardation.^23^ In ameloblasts, Wdr72 is thought to regulate vesicle trafficking and pH homeostasis via V-ATPase-dependent mechanisms.^23^ To specifically dissect the renal-specific function of Wdr72 we generated a kidney-specific knockout model.^38,39^ Several murine models of human dRTA illustrate a spectrum of dRTA phenotypes. Mice lacking *Atp6v0a4*^40^, *Slc4a1*^41^ or *Foxi1*^41,42^ develop severe systemic acidosis under basal conditions, whereas *Atp6v1b1* homozygous^34^ mutants as well as *Atp6v1b1*^35^ and *Atp6v0a4*^40^ heterozygotes, exhibit incomplete dRTA that becomes evident only upon acid challenge. Our model also exhibited features of incomplete distal RTA, characterized by impaired urinary acidification under both baseline and acid-loading conditions.

Our model exhibited hallmarks of incomplete dRTA, with persistently elevated urinary pH as well as reduced titratable acid excretion under both baseline and acid-loaded conditions. Although net acid excretion was lower at baseline, it normalized after chronic acid loading, likely reflecting activation of compensatory mechanisms. Notably, renal ammoniagenesis markers increased already under baseline conditions in *Wdr72*-deficient mice, even though urinary NH₄⁺ excretion was unchanged. Upon chronic acid challenge, urinary NH₄⁺ excretion and ammoniagenesis were both upregulated, indicating a robust compensatory mechanism^43^ that may account for the overall mild systemic acidosis. The elevated acid-base score, which integrates urinary pH and NH₄⁺ excretion to assess tubular acid excretion capacity,^29^ further suggests a potential proximal tubular adaptation to distal acidification defects.

The blood gas analysis in *Wdr72^fl/fl^;Pax8 Cre^+^*upon chronic acid load, is consistent with a mixed acid–base disorder. The overall low pH and relatively high pCO₂ observed across all groups (knockout vs. control in standard diet, acute and chronic acid load) likely reflects anesthesia-induced hypoventilation during terminal cardiac puncture, which was the only method approved by our local animal welfare authorities for blood collection. However, following chronic acid loading, *Wdr72^fl/fl^;Pax8 Cre^+^* mice exhibited mild hyperchloremic, hypokalemic acidosis - electrolyte alterations typical of incomplete dRTA which are consistent with observations in the *Atp6v1b1* knockout model.^35^ Importantly, these changes were not attributable to dehydration, as urine output, body weight, and hematocrit remained comparable between genotypes. Standardization of anesthetic exposure, sampling interval, and analysis minimized procedural variability in our experiments; nevertheless, the elevated pCO₂ may have partially masked the metabolic acidosis expected from impaired distal proton secretion, particularly under acid-loaded conditions. These findings underscore the need for alternative sampling strategies, such as arterial sampling in conscious or mechanically ventilated mice, to accurately assess systemic acid–base status.

Nephrocalcinosis and hypercalciuria, frequent features in patients with dRTA, were absent in 9 and 23 week old *Wdr72*-deficient mice. However, the possibility of late-onset phenotypes in older mice cannot be excluded. Instead, these animals displayed hypocalciuria upon baseline and acute acid load compared with controls, a finding also reported in *Atp6v1b1* and *Slc4a1* knock-in mouse models.^35,37,41^ The mechanisms underlying this paradoxical reduction in urinary calcium excretion remain unclear but may relate to the more alkaline urinary milieu or species-specific differences in tubular architecture, diet, or compensatory capacity. Skeletal manifestations were not assessed in our kidney-specific model, as the animals displayed normal electrolytes under standard diet. Overall, the phenotype of *Wdr72^fl/fl^;Pax8 Cre^+^*, contrasts with the early-onset and severe metabolic acidosis observed in patients harboring biallelic variants in *WDR72*, suggesting partial redundancy or compensation within the murine distal nephron.

In the CD of *Wdr72^fl/fl^;Pax8 Cre^+^* mice, type A ICs were more numerous and failed to expand further in response to chronic acid loading. In control animals, acid challenge normally induces expansion of type A ICs to enhance their functional capacity to increase proton secretion.^17,44^ Under sustained acid stress, the increased number of type A ICs in mice can occur through a combination of mechanisms, including transdifferentiation of non- A, non- B ICs into A ICs, limited proliferation of existing ICs, and, to a lesser extent, phenotypic conversion of AQP2^+^ precursor cells.^17,44,45^ Additionally, existing type A ICs can enhance their functional capacity by increasing apical membrane area and trafficking more V-ATPase to the apical surface.^17^ The impaired adaptive response in *Wdr72*-deficient mice suggests that WDR72 is critical for IC plasticity, enabling structural and functional remodeling of the distal nephron under acid stress.

In *Wdr72^fl/fl^;Pax8 Cre^+^* mice, type A ICs exhibit a reduced abundance and mislocalization of the B1 subunit, despite unaltered transcript levels, indicating a post-transcriptional defect that impairs V-ATPase stability, trafficking or assembly, and highlighting a central role for WDR72 in coordinating V-ATPase regulation. Notably, this phenotype parallels that of IC-specific *Dmxl1* knockout mice^36^, in which defective V-ATPase assembly similarly causes incomplete dRTA, highlighting a potential mechanistic convergence between WDR72 and DMXL1 in regulating acid–base handling in the distal nephron.

In higher eukaryotes, reversible V-ATPase assembly and trafficking are primarily regulated by the Rabconnectin-3 complex, which comprises two α subunits (DMXL1 and DMXL2) and a single β subunit, WDR7. Importantly, Rabconnectin-3 is not a uniform entity; distinct combinations of DMXL isoforms with WDR7 generate tissue- and compartment-specific complexes that selectively regulate subsets of V-ATPases. For example, DMXL2/WDR7 complexes in neurons facilitate high-demand synaptic vesicle acidification, ensuring efficient neurotransmitter loading, while in non-neuronal cells, DMXL1/WDR7 complexes regulate V-ATPase assembly.^46–48^ This modular organization allows Rabconnectin-3 complexes to fine-tune V-ATPase activity according to cellular context and physiological demand.

Mammals express a highly conserved homolog of WDR7, WDR72.^21^ WDR72 is specifically expressed at high levels in the kidney.^22^ Although direct interactions with canonical Rabconnectin-3 subunits have not yet been demonstrated, its structural similarity to WDR7, enrichment in ICs, and association with dRTA suggest a potential role in V-ATPase regulation in a cell type–specific manner. *Wdr72^fl/fl^;Pax8 Cre^+^* mice show reduced V-ATPase B1 subunit abundance and membranous targeting, consistent with impaired proton secretion, and reminiscent of phenotypes reported for IC-specific *Dmxl1* knockout mice, where defective V-ATPase assembly leads to incomplete dRTA.

Our findings establish *Wdr72* as a critical modulator of distal nephron function *in vivo*. Our data implicate Wdr72 in V-ATPase stabilization, assembly or trafficking, and suggest mechanistic insight into how its deficiency might contribute to distal acidification defects in humans. Future studies will be essential to define WDR72’s interactions within Rabconnectin-3 complexes and to refine approaches for capturing systemic acid–base responses under physiologic conditions.

## Acknowledgments

We gratefully acknowledge Dr. Dülsner and FEM-CVK for supporting adherence to the 3R principles within our local mouse facilities. We thank Kerstin Sommer and Sandra Gerstenberg for technical support in the genotyping process.

## Author contributions

Conceptualization: V.K., A.A.S.

Formal analysis: A.A.S., P.M., T.B., M.K., S.W

Funding acquisition: V.K.

Methodology: V.K., D.M., B.P., P.M., T.B., A.A.S.

Project administration: V.K.

Resources: V.K., D.M., B.P., M.K

Supervision: V.K., D.M., B.P.

Validation: V.K., A.A.S., P.M., M.K.

Visualization: V.K., A.A.S., P.M.

Writing – original draft: V.K., A.A.S., P.M.

Writing – review & editing: V.K., A.A.S., P.M.

## Sharing Statement

All data will be shared upon request.

## Supplemental material

**Supplemental Table 1. Summary of body measurements of *Wdr72^fl/fl^;Pax8 Cre^-^* (Cre-) vs. *Wdr72^fl/fl^;Pax8 Cre^+^* (Cre+) mice collected under standard diet, acute and chronic acid load conditions.**

**Supplemental Table 2. Comparison of blood gas analysis of control mice under standard diet, acute and chronic acid load condition**

**Supplemental Figure 1. Standard curve validation**

**Supplemental Figure 2. Identification of 21 variants in *WDR72* in 26 families with dRTA and AI.**

**Supplemental Figure 3. W*d*r72 expression across nephron cell types based on single-cell RNA sequencing datasets.**

**Supplemental Figure 4. Loss of *Wdr72* does not cause renal nephrocalcinosis.**

**Supplemental Figure 5. Heatmap showing mRNA expression levels of different subunits of V-ATPases, intercalated cell (IC) marker genes, and ammoniagenesis-related genes.**

**Supplemental Figure 6. Wdr72 does not influence the PC:IC pattern under acid conditions.**

**Supplemental Figure 7. Subcellular distribution of Atp6v1b1 in *Wdr72-deficient* kidneys.**

