## Supplemental Material for "Kidney-specific *Wdr72* deletion leads to incomplete distal renal tubular acidosis through impaired V-ATPase B1 subunit localization"

**Supplemental Table 1. Summary of body measurements of *Wdr72$^{fl/fl}$;Pax8 Cre$^{-}$* (Cre-) vs. *Wdr72$^{fl/fl}$;Pax8 Cre$^{+}$* (Cre+) mice collected under standard diet, acute and chronic acid load conditions.**
**Supplemental Table 2. Comparison of blood gas analysis of control mice under standard diet, acute and chronic acid load condition**

**Supplemental Figure 1. Standard curve validation**
**Supplemental Figure 2. Identification of 21 variants in *WDR72* in 26 families with dRTA and AI.**
**Supplemental Figure 3. *Wdr72* expression across nephron cell types based on single-cell RNA sequencing datasets.**
**Supplemental Figure 4. Loss of *Wdr72* does not cause renal nephrocalcinosis.**
**Supplemental Figure 5. Heatmap showing mRNA expression levels of different subunits of V-ATPases, intercalated cell (IC) marker genes, and ammoniagenesis-related genes.**
**Supplemental Figure 6. Wdr72 does not influence the PC:IC pattern under acid conditions.**
**Supplemental Figure 7. Subcellular distribution of Atp6v1b1 in *Wdr72-deficient* kidneys.**

| Parameter (unit) | Standard diet | | | | | Acute acid load | | | | | Chronic acid load | | | | |
|---|---|---|---|---|---|---|---|---|---|---|---|---|---|---|---|
| | Cre- | | Cre+ | | p-Value | Cre- | | Cre+ | | p-Value | Cre- | | Cre+ | | p-Value |
| | mean | ± SEM | mean | ± SEM | | mean | ± SEM | mean | ± SEM | | mean | ± SEM | mean | ± SEM | |
| Urine volume (ml) | 7.43 | 1.04 | 5.85 | 0.89 | 0.30 | 2.16 | 0.10 | 2.23 | 0.17 | 0.74 | 1.99 | 0.18 | 1.66 | 0.11 | 0.18 |
| Water consumption (ml) | 10.10 | 1.69 | 8.28 | 0.90 | 0.39 | 2.70 | 0.24 | 2.72 | 0.12 | 0.95 | 2.43 | 0.12 | 2.48 | 0.09 | 0.77 |
| Feces (g) | 0.41 | 0.05 | 0.45 | 0.05 | 0.54 | 0.32 | 0.03 | 0.38 | 0.02 | 0.12 | 0.38 | 0.03 | 0.53 | 0.07 | 0.09 |
| Food consumption (g) | 3.13 | 0.15 | 3.18 | 0.24 | 0.88 | 2.79 | 0.12 | 3.11 | 0.20 | 0.22 | 2.92 | 0.13 | 2.94 | 0.11 | 0.91 |
| Weight (g) | 27.31 | 0.45 | 25.33 | 0.93 | 0.10 | 26.33 | 0.41 | 25.76 | 0.76 | 0.56 | 28.04 | 0.29 | 27.07 | 0.56 | 0.19 |
| Weight loss in metabolic cages (g) | 2.36 | 0.15 | 1.98 | 0.31 | 0.33 | 2.47 | 0.22 | 2.54 | 0.21 | 0.84 | 2.53 | 0.24 | 2.26 | 0.13 | 0.38 |
| Weight loss in 6 days acid load (g) | NA | NA | NA | NA | NA | NA | NA | NA | NA | NA | 0.42 | 0.34 | 1.20 | 0.22 | 0.10 |

**Supplemental Table 1. Summary of body measurements of *Wdr72$^{fl/fl}$;Pax8 Cre$^{-}$* (Cre-) vs. *Wdr72$^{fl/fl}$;Pax8 Cre$^{+}$* (Cre+) mice collected under standard diet, acute and chronic acid load conditions.** Statistical analyses were performed using unpaired t-tests; values are reported as means +/- SEM; p-values <0.05 were considered significant; n=7-8. g, gram; mL, milliliter.

| | Standard diet | | Acute acid load | | Chronic acid load | | Standard diet vs. acute acid load | Standard diet vs. chronic acid load | Acute acid load vs. chronic acid load |
|---|---|---|---|---|---|---|---|---|---|
| | Cre- | | Cre- | | Cre- | | | | |
| | mean | ± SEM | mean | ± SEM | mean | ± SEM | | | |
| Blood pH | 7.20 | 0.01 | 7.10 | 0.03 | 7.20 | 0.02 | 0.01 | 0.92 | 0.01 |
| $pCO_2$ (mmHg) | 69.19 | 1.75 | 63.46 | 1.43 | 67.36 | 1.60 | 0.09 | 0.46 | 0.26 |
| $cHCO_3^-(P)c$ (mM) | 25.53 | 1.04 | 19.11 | 1.21 | 25.40 | 1.18 | 0.004 | 0.94 | 0.004 |
| $cHCO_3^-(Pst)c$ (mM) | 19.43 | 0.83 | 14.31 | 1.09 | 19.51 | 0.94 | 0.01 | 0.95 | 0.01 |
| ABE*c* (mM) | -4.11 | 1.13 | -11.70 | 1.70 | -3.97 | 1.29 | 0.005 | 0.95 | 0.005 |
| SBE*c* (mM) | -1.73 | 1.11 | -9.07 | 1.51 | -1.73 | 1.33 | 0.004 | 1.00 | 0.004 |
| Hct*c* (%) | 44.44 | 0.41 | 44.30 | 0.61 | 43.20 | 0.81 | 0.88 | 0.52 | 0.52 |
| CRE (mM) | 0.01 | 0.00 | 0.01 | 0.00 | 0.01 | 0.00 | 0.22 | 0.45 | 0.52 |
| Urine pH | 6.82 | 0.04 | 6.54 | 0.06 | 5.87 | 0.04 | 0.001 | <0,001 | <0,001 |
| AB score | 23.31 | 1.79 | 28.92 | 0.90 | 26.29 | 0.56 | 0.02 | 0.22 | 0.22 |
| NAE (umol/h) | 4.21 | 0.41 | 4.60 | 0.52 | 10.96 | 1.18 | 0.76 | <0,001 | <0,001 |
| $[TA - HCO_3^-]$/CRE | 3.08 | 1.54 | 11.34 | 2.50 | 15.41 | 0.75 | 0.01 | <0,001 | 0.13 |
| $NH_4^+$ /CRE | 12.42 | 0.52 | 18.61 | 1.47 | 35.64 | 3.29 | 0.11 | <0,001 | <0,001 |

**Supplemental Table 2. Comparison of blood gas analysis of control mice under standard diet, acute and chronic acid load condition.** Statistical analyses were performed using one-way ANOVA with Holm Sidak's test; values are reported as means +/- SEM; p-values <0.05 were considered significant; n=6-7. C, calculated; ABE, actual base exess; SBE, standard base exess; Hct, hematocrit; CRE, createnine; AB score, acid-base score; NAE, net acid excretion; TA, titratable acids.

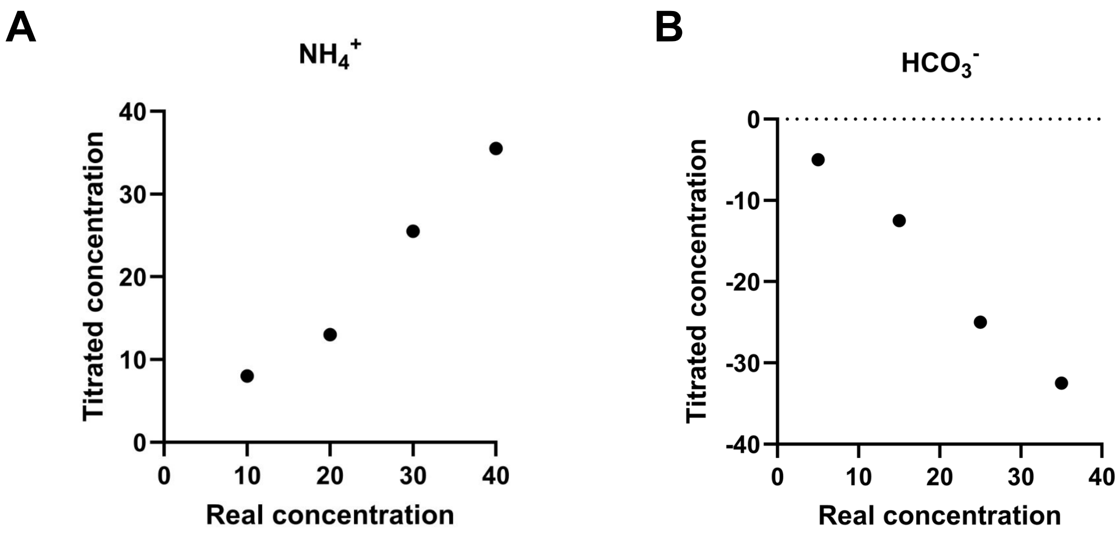


**Supplemental Figure 1. Standard curve validation** of the measurement of ammonium (**A**), and bicarbonate ($HCO_3^-$) (**B**) using the titrimetric method conducted in this study.

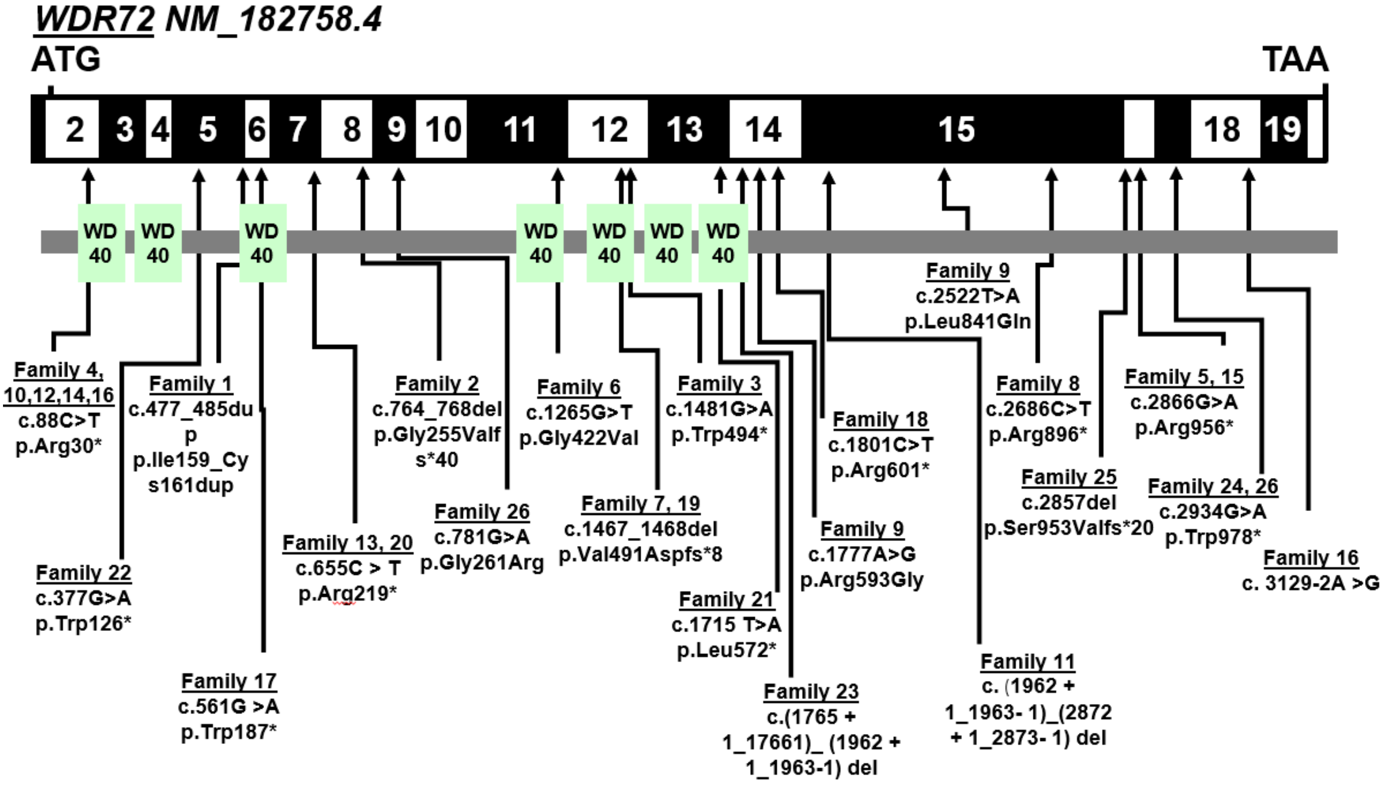


**Supplemental Figure 2. Identification of 21 variants in *WDR72* in 26 families with dRTA and AI.** Exon structure and protein domains of WDR72 are shown. Position of start codon (ATG) and stop codon (TAA) are indicated. Numbers indicate different exons, which are marked by black or white boxes. WD40 domains are shown by colored boxes below. *WDR72* variant positions are indicated by a black arrow in relation to the exon and protein domain. Variants in family 1 and 2 were discovered by us (Jobst-Schwan, Klämbt *et al.*, 2020). References for family 3, 4 and 5 (Khandelwal *et al.*, 2021), family 6 and family 7 (Zhang et al., 2019) and 9 (Rungroj *et al.*, 2018), 10- 24 (Deepthi et al., 2025), 25 (Al-Omairi, et al., 2025), and 26 (Gupta et al, 2025). Note that family 7 and 24 has a compound heterozygous mutation, while all other families have homozygous variants. Note that mutations in family 7, 18 and 24 had previously been described in cases of amelogenesis imperfecta but were novel in cases of dRTA.

**A** Healthy Mouse Dataset: Wu et al, JASN 2019; and RBK RID: 14-4KBC.

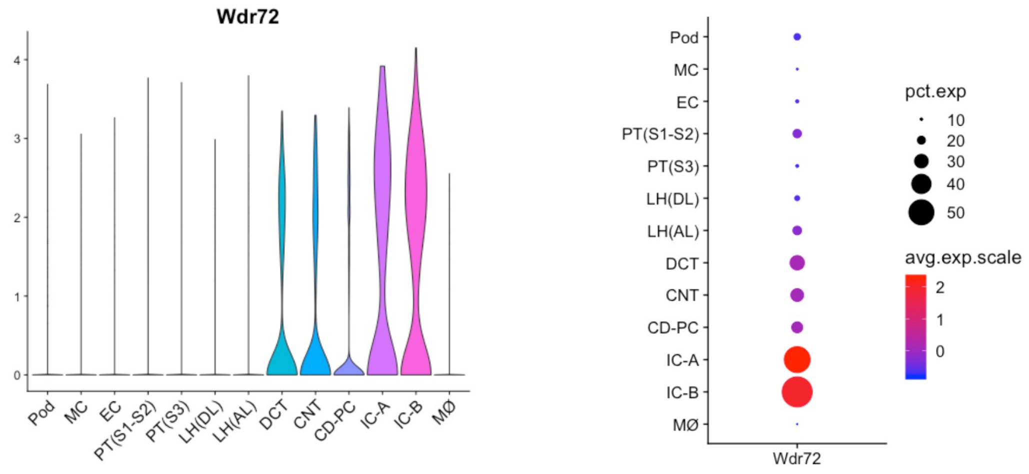


**B**

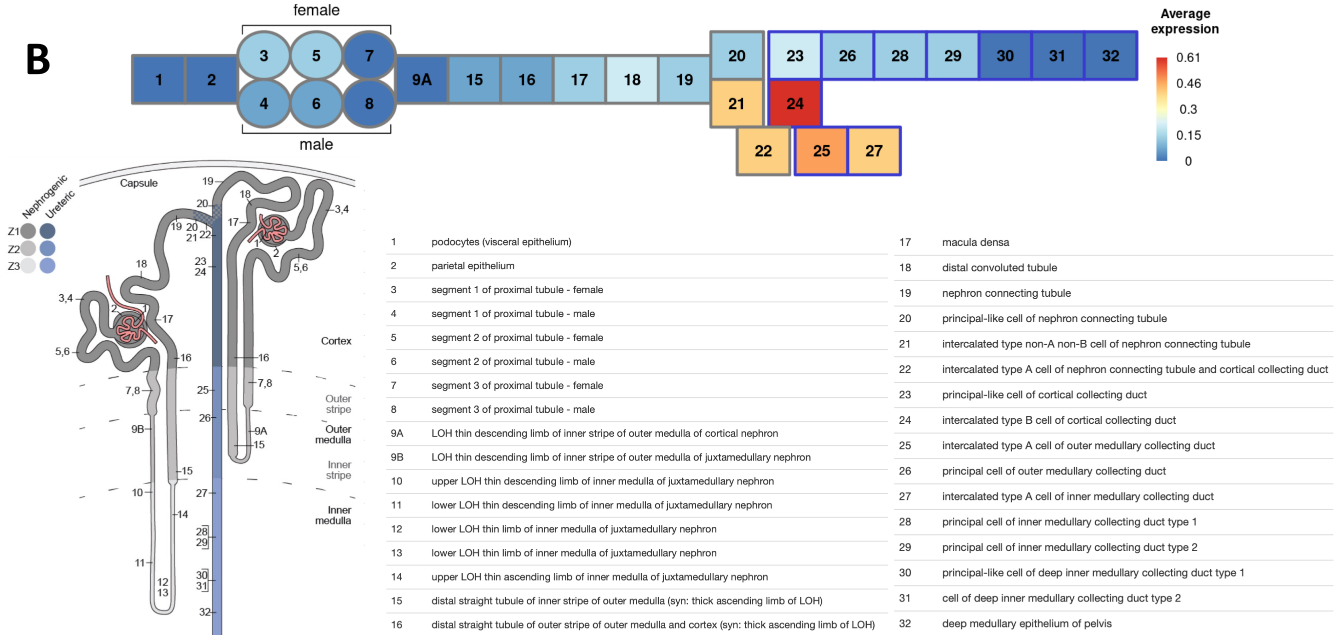


**Supplemental Figure 3. *Wdr72* expression across nephron cell types based on single-cell RNA sequencing datasets.** (**A**) *Wdr72* is highly expressed in type A ICs, type B ICs (CD), and to a lesser extend in the PC (CD), DCT, and CNT in mouse kidney (Wu et al. 2019., https://humphreyslab.com/SingleCell/) (**B**) *Wdr72* is highly expressed in type B ICs (CCD), type A ICs (CNT, CCD, OMCD, and IMCD), non A- non- B IC (CNT) (Ransick et al. 2019, https://cello.shinyapps.io/kidneycellexplorer/). CD-PC, collecting duct principal cells; CNT, connecting tubule; DCT, distal convoluted tubule; EC, endothelial cells; IC-A, type A intercalated cells; IC-B, type B intercalated cells; LH, loop of Henle; M, macrophages; MC, mesangial cells; Pod, podocytes; PT, proximal tubule (S1–S3 segments).

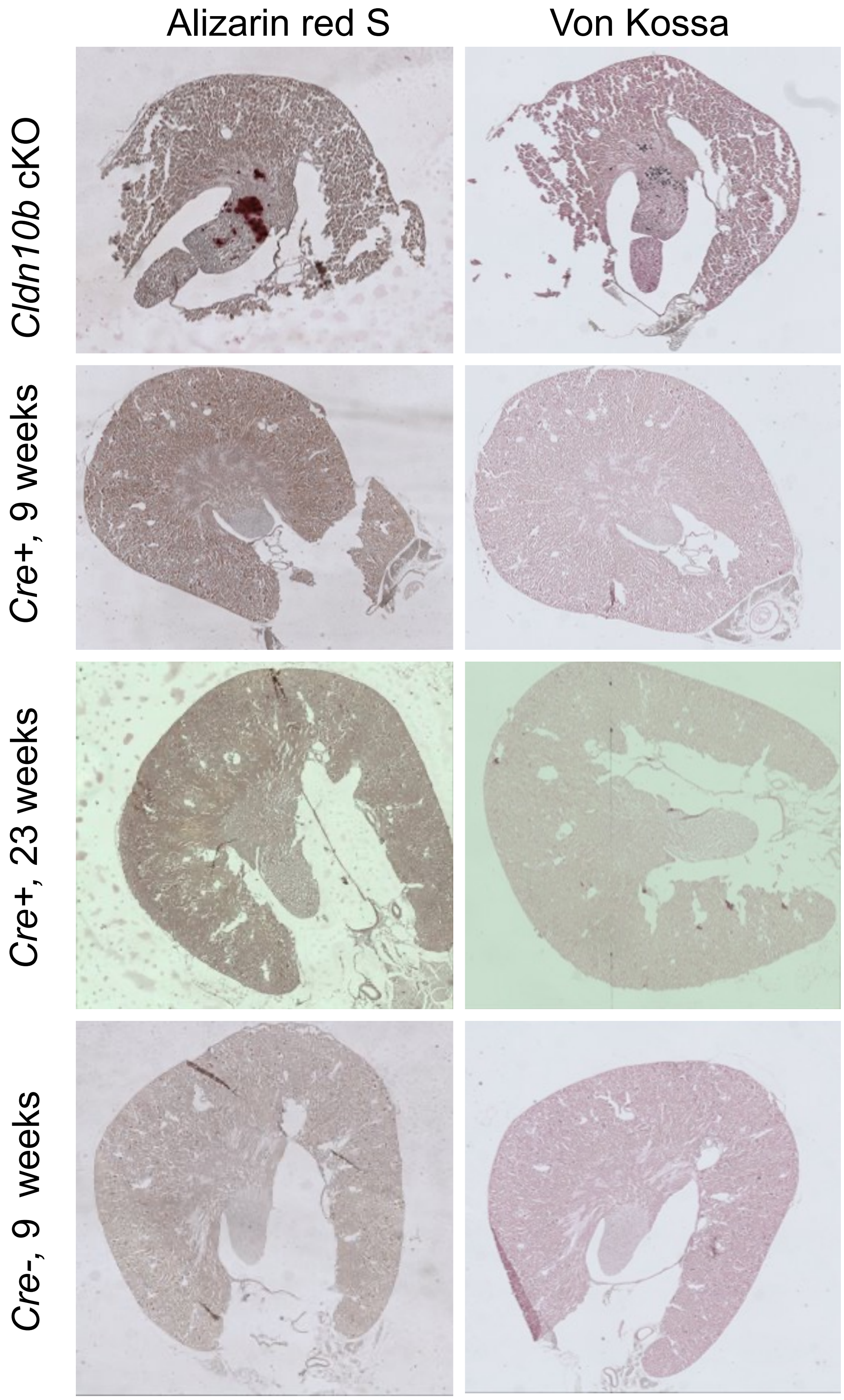


**Supplemental Figure 4. Loss of *Wdr72* does not cause renal nephrocalcinosis.** To evaluate renal calcifications, kidneys from *Wdr72*$^{fl/fl}$*;Pax8 Cre*$^{+}$ (Cre+) vs. *Wdr72*$^{fl/fl}$*;Pax8 Cre*$^{-}$ (Cre-) mice were analyzed at 9 and 23 weeks of age using Alizarin Red S (calcium deposit) and von Kossa (phosphate/carbonate deposits) staining. *Cldn10b* cKO mice were stained in parallel as positive controls. For each group, *n*=3 mice were examined. Staining patterns were evaluate qualitatively from stitched 10× composite images. Note: *Wdr72*$^{fl/fl}$*;Pax8 Cre*$^{+}$ (Cre+) vs. *Wdr72*$^{fl/fl}$*;Pax8 Cre*$^{-}$ (Cre-) showed no evidence of nephrocalcinosis at neither 9 nor 23 weeks.

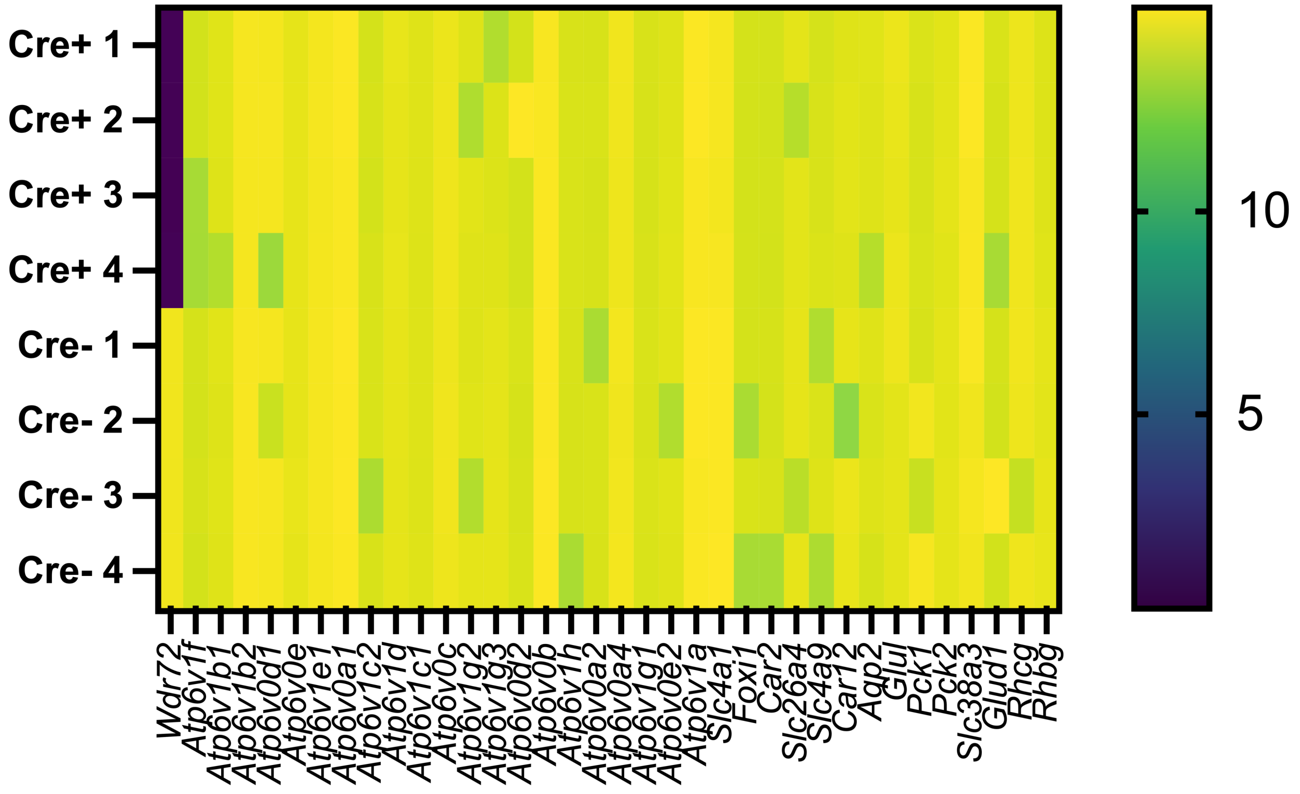


**Supplemental Figure 5. Heatmap showing mRNA expression levels of different subunits of V-ATPases, intercalated cell (IC) marker genes, and ammoniagenesis-related genes.** Bulk RNA-sequencing was performed on kidneys from 4 *Wdr72*$^{fl/fl}$*;Pax8 Cre+* and 4 *Wdr72*$^{fl/fl}$*;Pax8 Cre-* mice that were fed standard diet. Expression values are shown as log(FPKM + 1). Each column represents a gene, and each row represents an individual sample. Color intensity corresponds to relative expression levels as indicated by the scale bar. Note: *Wdr72* expression is altered in Wdr72$^{fl/fl}$;*Pax8 Cre+* kidneys, indicating the knockout, whereas other genes show no differential expression under standard diet conditions.

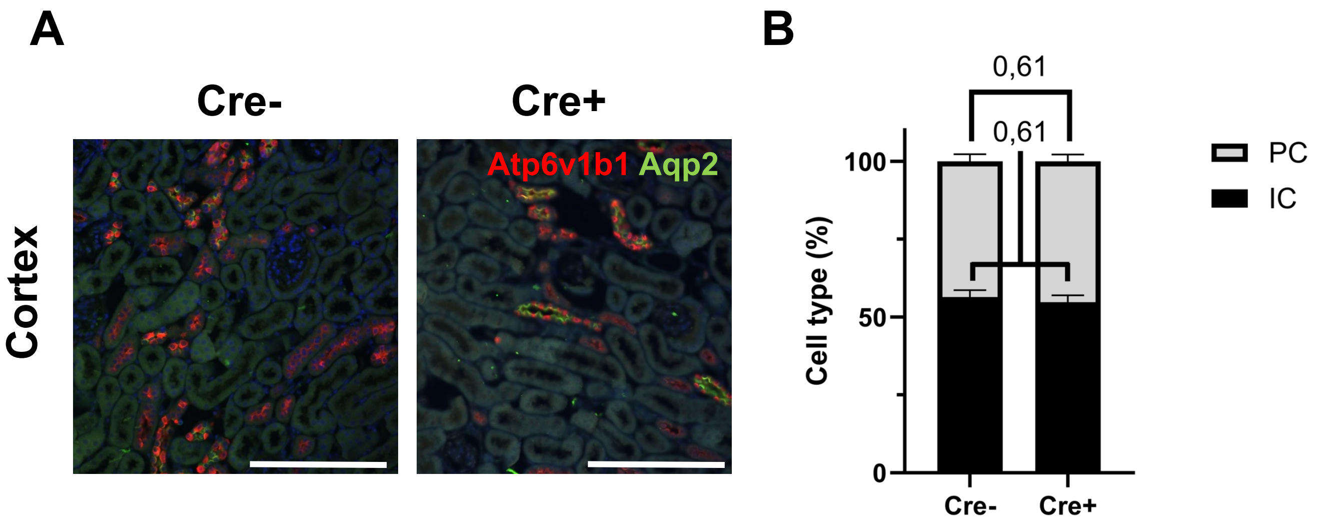


**Supplemental Figure 6. Wdr72 does not influence the PC:IC pattern under acid conditions.** (**A**) Representative immunofluorescence images of cortical collecting duct cells under chronic acid load. Kidneys were stained for Aqp2 (green) and Atp6v1b1 (red); nuclei were counterstained with DAPI (blue). Aqp2-positive cells were identified as PCs and Atp6v1b1-positive cells as ICs. (**B**) Under chronic acid load, the IC-to-PC ratio remained unchanged between genotypes in cortical CD. Statistical comparisons were performed using 2way ANOVA; values are reported as means +/- SEM; $p < 0.05$ was considered significant; n=3. Scale bar=200 µm.

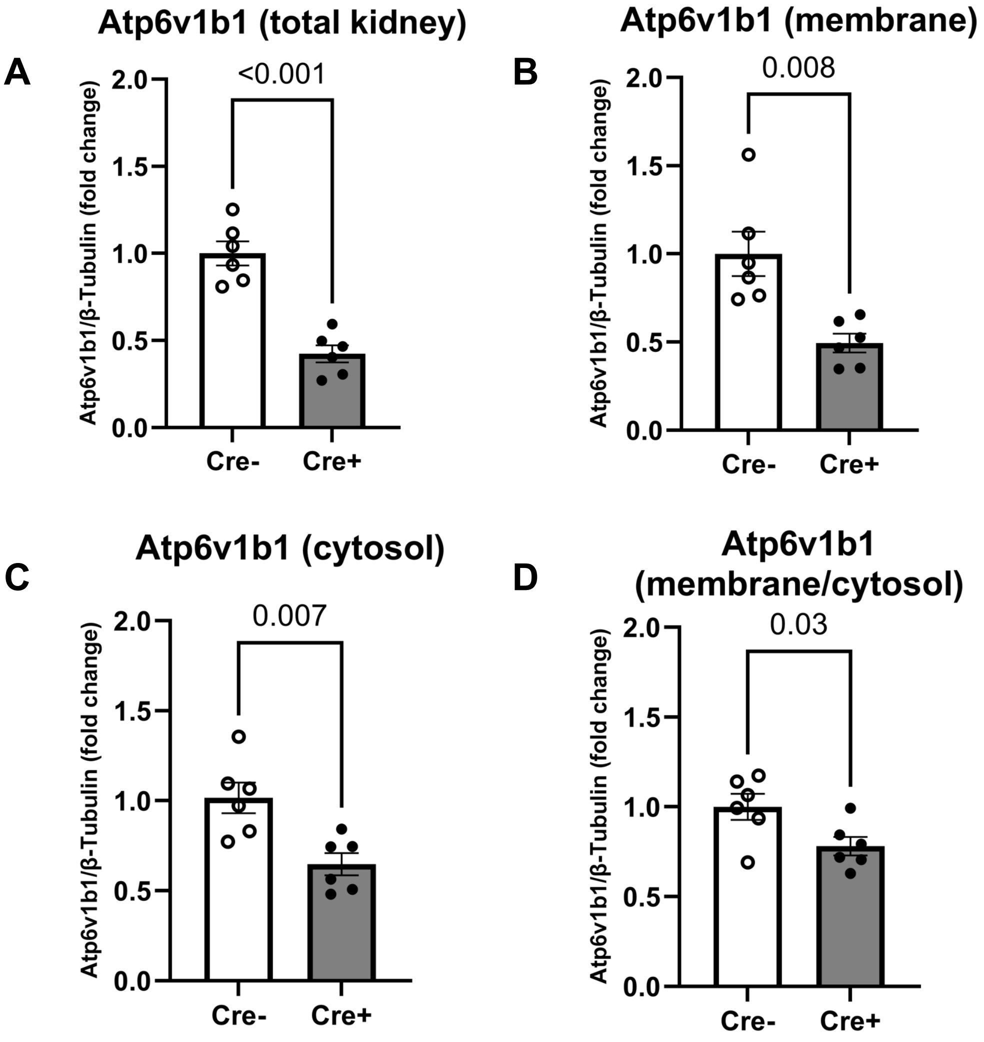


**Supplemental Figure 7. Subcellular distribution of Atp6v1b1 in *Wdr72-deficient* kidneys. (A–D**) Quantification of Western blot analyses corresponding to Figure 6E from subcellular fractionation of kidneys from *Wdr72*$^{fl/fl}$*;Pax8 Cre*$^{+}$ (Cre+) vs. *Wdr72*$^{fl/fl}$*;Pax8 Cre*$^{-}$ (Cre-) mice. The abundance of the Atp6v1b1 was reduced in total lysates (**A**), the membrane fraction (**B**), and the cytosolic fraction (**C**) in Cre$^{+}$ kidneys. (**D**) The membrane-to-cytosol ratio of Atp6b1b1 was also decreased, indicating impaired subcellular distribution. Protein levels were normalized to β-tubulin. Statistical comparisons were performed using unpaired t-tests; values are reported as means +/- SEM; $p < 0.05$ was considered significant; n=6.
